# A tale of caution: How endogenous viral elements affect virus discovery in transcriptomic data

**DOI:** 10.1101/2023.09.08.556789

**Authors:** Nadja Brait, Thomas Hackl, Côme Morel, Antoni Exbrayat, Serafín Gutierrez, Sebastian Lequime

## Abstract

Large-scale metagenomic and -transcriptomic studies have revolutionized our understanding of viral diversity and abundance. In contrast, endogenous viral elements (EVEs), remnants of viral sequences integrated into host genomes, have received limited attention in the context of virus discovery, especially in RNA-Seq data. EVEs resemble their original viruses, a challenge that makes distinguishing between active infections and integrated remnants difficult, affecting virus classification and biases downstream analyses. Here, we systematically assess the effects of EVEs on a prototypical virus discovery pipeline, evaluate their impact on data integrity and classification accuracy, and provide some recommendations for better practices.

We examined EVEs and exogenous viral sequences linked to Orthomyxoviridae, a diverse family of negative-sense segmented RNA viruses, in 13 genomic and 538 transcriptomic datasets of Culicinae mosquitoes. Our analysis revealed a substantial number of viral sequences in transcriptomic datasets. However, a significant portion appeared not to be exogenous viruses but transcripts derived from EVEs. Distinguishing between transcribed EVEs or exogenous virus sequences was especially difficult in samples with low viral abundance. For example, three transcribed EVEs showed full-length segments, devoid of frameshift and nonsense mutations, exhibiting sufficient mean read depths that qualify them as exogenous virus hits. Mapping reads on a host genome containing EVEs before assembly somewhat alleviated the EVE burden, but it led to a drastic reduction of viral hits and reduced quality of assemblies, especially in regions of the viral genome relatively similar to EVEs.

Our study highlights that our knowledge of the genetic diversity of viruses can be altered by the underestimated presence of EVEs in transcriptomic datasets, leading to false positives and altered or missing sequence information. Thus, recognizing and addressing the influence of EVEs in virus discovery pipelines will be key to enhancing our ability to capture the full spectrum of viral diversity.

## Introduction

Decreasing costs of high throughput sequencing has allowed viral metagenomics to grow exponentially, with skyrocketing numbers of viral species deposited yearly (Wolf et al., 2020). Consequently, computationally generated genome assemblies have become sufficient evidence for their admission as *bona fide* viruses, no longer relying on cultivation and verification through phenotypic properties (Simmonds et al., 2017). A substantial subset of deposited genomes is derived from the re-analysis of genomic or transcriptomic datasets that were not always intended for virus discovery (e.g., Edgar et al., 2022; Johansen et al., 2022; Nayfach et al., 2021; Neri et al., 2022; Shi et al., 2016; Simmonds et al., 2017). While this opens up the spectrum of associated hosts and expands our knowledge of virus diversity, contaminating host nucleic acids are usually not removed before sequencing, taking away sequence capacity for viral reads from an already potentially low viral load in the samples (Prachayangprecha et al., 2014). The low abundance of viral reads combined with the presence of related species or populations in the same sample makes viral assembly challenging. Viruses with low and uneven read depth are known to produce fragmented assemblies or chimeric contigs if they are similar enough (García-López et al., 2015; Smits et al., 2015; Sutton et al., 2019). Viral reads with high sequence similarity may not only originate from similar viral populations in the host but could potentially come from the host genome itself in the form of prophages, provirus, or (non-) retroviral integration (Ackermann, H. W., and M. S. DuBow., 1987; Weiss, 2006; Zhdanov, 1975).

Endogenous viral elements (EVEs) are partial or complete insertions of viral genomes in the host’s genome (Bejarano et al., 1996; Benveniste & Todaro, 1974; Crochu et al., 2004; Jaenisch, 1976). For such endogenization to occur, two essential steps need to happen: (I) the production of dsDNA intermediates and (II) the integration into the host germline, probably through nonhomologous recombinations or reverse transcription and integration driven by cellular retroelements (Belyi et al., 2010; Geuking et al., 2009; Holmes, 2011; Tassetto et al., 2019; Wallau, 2022). Most EVEs are of retroviral origin, as the integration of retroviruses is an obligatory part of their replication cycle (Herniou et al., 1998; Katzourakis et al., 2005). Retroviral EVEs were first described in the 1970s (Benveniste & Todaro, 1974), and despite non-retroviral EVEs being experimentally induced in the same decade (Zhdanov, 1975), the first non-retroviral EVE in metazoans was only described 30 years later (Crochu et al., 2004). Since then, all Baltimore classes of viruses have been found to be integrated (Berns & Linden, 1995; Geisler & Jarvis, 2016; Horie et al., 2010; Katzourakis et al., 2007; H. Liu et al., 2011; W. Liu et al., 2012; Staginnus & Richert-Pöggeler, 2006; Taylor et al., 2010).

Recent insertions and purifying selections within the host genome can lead to the emergence of endogenous viral elements (EVEs) similar to currently circulating viruses (Aiewsakun & Katzourakis, 2015). The misclassification of EVE sequences as exogenous viruses is a particularly underestimated possibility in virus discovery studies based on (meta)-transcriptomics datasets. Indeed, depending on their integration sites, EVEs can exploit nearby regulatory elements and the host cell’s transcriptional machinery, facilitating EVE transcription (Sofuku et al., 2018). While most described EVEs are highly mutated or fragmented, transcribing as non-coding RNAs, several have been discovered to encode intact open reading frames (ORFs) (Horie et al., 2010; Katzourakis & Gifford, 2010). In addition, during viral genome assembly, EVE-associated reads exhibiting high sequence similarity to exogenous viruses can inadvertently integrate into chimeric viral contigs or increase assembly fragmentation. Consequently, the accurate reconstruction and characterization of complete viral genomes becomes challenging. The common practice of host read removal in virus discovery pipelines, which could alleviate the burden of EVEs, can unintentionally exclude valuable viral-associated reads along with EVE sequences.

In this study, we aimed to examine the potential bias of transcribed EVEs in interpreting viral sequence assemblies in typical virus discovery pipelines. We focused our study on orthomyxoviruses in Culicinae mosquitoes. Orthomyxoviridae is a family of enveloped segmented negative-sense single-stranded RNA viruses that mainly infect vertebrates and arthropods. Since the first description of influenza A virus in 1933, *Orthomyxovirus* discovery has been primarily driven by public health risk, comprising only a small group of mammal, bird, and tick-associated RNA viruses (Allison et al., 2015; Presti et al., 2009), but recent metatranscriptomics studies have detected a vast amount of novel orthomyxoviruses in invertebrates, greatly expanding the known host range and diversity of this family (Batson et al., 2021; Li et al., 2015; M. Shi et al., 2016). Complete orthomyxovirus genomes comprise between 6 and 8 segments, with some segments only recently being identified, as can be seen for the genus *Quaranjavirus,* with two additional hypothetical proteins discovered (Batson et al., 2021). Despite the recent increase in *Orthomyxovirus* diversity, most non-influenza genomes are not fully characterized, resulting in many species being taxonomically unclassified (Benson et al., 2013). Segmented viruses are often prone to loss of genomic information since most metagenomic studies predominantly screen only for the RNA-dependent RNA polymerase (RdRp). While non-segmented genomes can often be connected through linkage analyses, segmented viruses are more difficult to assemble, especially from diverse populations or low viral abundance (Krishnamurthy & Wang, 2017).

Our analysis of publicly available genomic and transcriptomic datasets from Culicinae mosquitoes revealed the presence of orthomyxovirus-derived EVEs integrated into host genomes, particularly in *Aedes* species. We show that classifying contigs into EVE transcripts or exogenous associated sequences for samples with low viral abundance is challenging and can potentially lead to misidentification. We also observed that EVEs in reference genomes for read removal can impact the downstream analysis of a typical virus discovery pipeline.

## Material and Methods

### Detection of EVEs in genomic datasets

Thirteen assemblies of whole genome sequence datasets of Culicinae mosquitoes were obtained from https://www.ncbi.nlm.nih.gov/Traces/wgs/ (Supplementary Table 1). Reference genomes, as well as transcript sequences from *Aedes albopictus* Foshan FPA, *Aedes aegypti* LVP_AGWG, and *Culex quinquefasciatus* Johannesburg, for host read removal were obtained from Vectorbase (Amos et al., 2022; Arensburger et al., 2010; Matthews et al., 2018; Palatini et al., 2020). Potential EVEs were screened and validated with an in-house R script (Figure 1, Scripts available on GitHub). To summarize, files were downloaded, merged, unzipped, and used as a database for a tblastn search (NCBI-blast-2.13.0) against curated *Orthomyxovirus* reference sequences (Accessions provided in Supplementary Table 2, sequences in Supplementary Data) with an e-value threshold of 10^-4^. Putative EVE sequences were extracted, clustered, and merged as long as they overlapped or had a gap size of less than 100 nucleotides and were in the same orientation on the host contig. Sequence clusters >= 250 nt were used as the queries for a reciprocal blast search (NCBI-blast-2.13.0) against the NCBI nr and nt databases with an e-value cut-off of 10^-4^ to ensure viral origin. Contigs of interest were extracted, and genetic features of identified EVEs were checked with the NCBI Conserved Domain Database (Marchler-Bauer et al., 2015). Amino acid sequences were obtained by translating query sequences with diamond blastx (v2.0.13.151) and were manually checked afterward. To account for potential frameshift mutations, alignments produced by diamond BLAST were screened with *blast traceback operations (BTOP)*. EVEs within the reference sequences were cut out manually.

**Figure 1:**
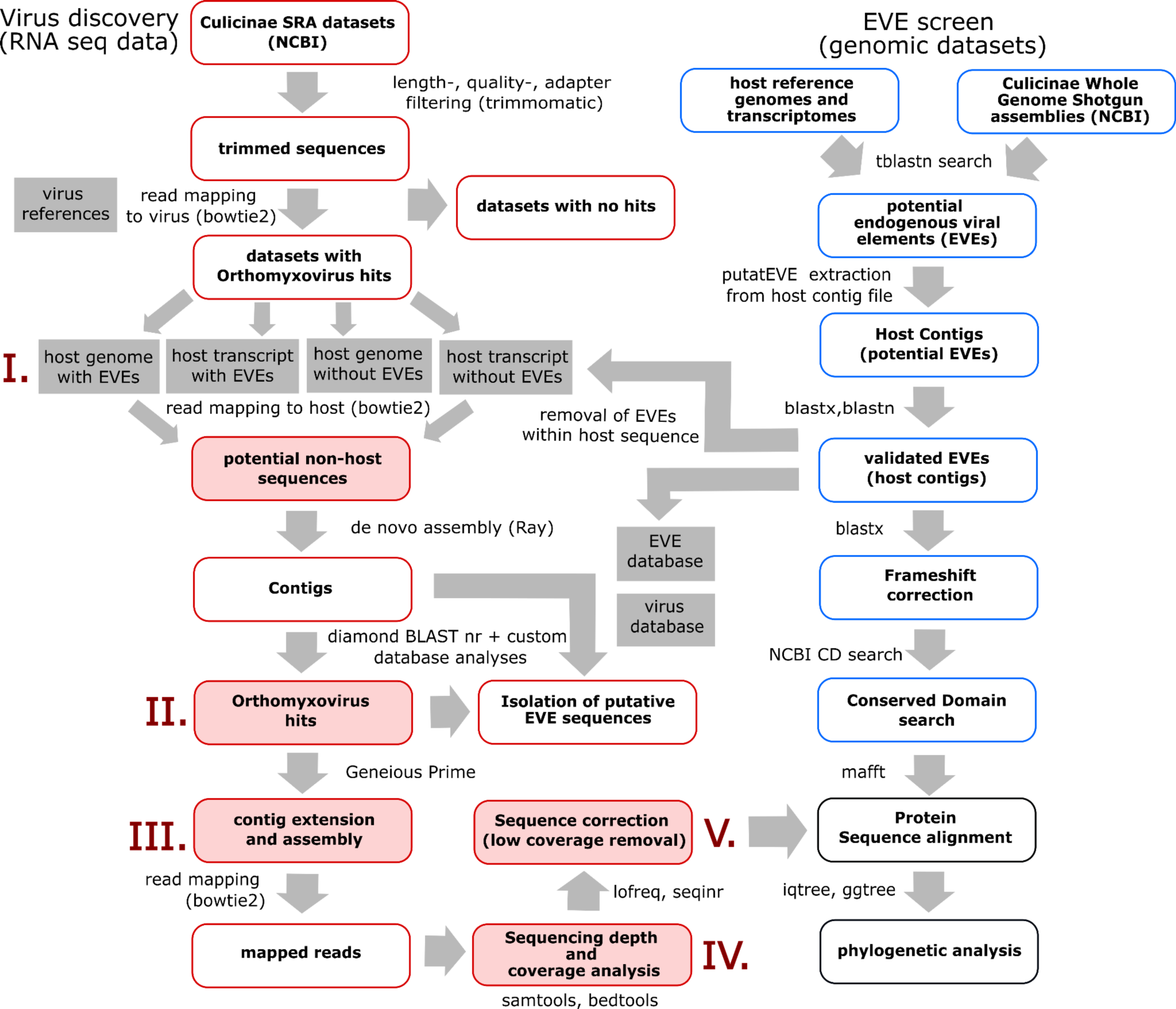
The influence of EVEs on the virus discovery process. Bioinformatics pipelines for transcriptomic virus discovery (tailored to exogenous orthomyxoviruses, framed in red) and genomic EVE detection (framed in blue). Pipeline steps potentially influenced by EVEs are shaded red: I. The presence of EVEs in host reference sequences may impact read filtering during host read removal. II. EVE sequences with high similarity to exogenous viruses could lead to false positive hits. III. EVE sequences with high similarity to exogenous viruses may erroneously assemble with exogenous viral sequences, resulting in chimeric assemblies. IV. Viral sequences accidentally filtered out during host read removal due to the presence of EVEs can lead to reduced sequencing depth and incomplete coverage. V. EVE sequences with high similarity to exogenous viruses might introduce SNPs in viral consensus sequences. The tools and packages utilized are indicated outside the respective boxes.

### Collection of metadata in the SRA repository

To identify datasets of high-throughput RNA-Seq of Culicinae, we searched the Sequence Read Archive (SRA) at NCBI (Leinonen et al., 2011). Datasets were sent to the SRA Run selector using the following search terms: “Aedes” [All Fields] OR “Culex” [All Fields] OR “Culicinae” [All Fields]). We excluded genomic datasets, amplicons, target capture, heavily modified clones, small RNAs (e.g., microRNA), and datasets with unspecified experimental workflows (Supplementary Table 6). Origins of mosquito latitude and longitude positions were approximated from metadata and plotted using ggplot2 (v3.3.5, (Whickham, 2016)) in the RStudio environment (v2021.09.1).

### Sequence assembly using publicly available transcriptomic data

Fastq files of RNA-Seq datasets were obtained from the SRA database by efetch (v14.6) and pre-processed using fastq-dump (v2.10.9). Sequences were checked with fastqc (v0.11.9), and adapter-, length- and quality trimmed with trimmomatic (v0.39). The trimmed reads were then passed to a relaxed Bowtie2 mapping with an *Orthomyxovirus*-specific reference list (Supplementary Table 2). Host read removal was carried out by mapping the trimmed sequences of samples with putative positive hits relaxed against the above-mentioned host genomes and transcriptomes. This step was performed for the references with still integrated EVEs and those with initial EVE removal. Unmapped reads were *de novo* assembled with Ray (version 2.3.1). Assembled contigs were screened against the nr database with a full sensitivity for hits of >40% identity from diamond-blastx (v2.0.13.151). Software, versions and parameters can be found in the Supplementary Table under STAR methods.

### Exploration of putative EVE-like sequences in transcriptomic data

We generated two custom databases of the existing *Orthomyxovirus* reference list or the translated amino-acid protein sequences of previously detected genomic EVEs. The highest bit score hits of viral contigs against each database were merged into a table, and a Δbit-score value (bit score values of exogenous virus hits – bit score values of EVE hits) was determined. Bit scores to EVE hits were assigned a negative value, while hits corresponding to the non-EVE references remained positive. Pmax values (parallel maxima of two vectors, consisting of either the highest EVE or non-EVE bit score) and Δbit-scores were plotted with geom_jitter (ggplot2 v.3.3.5). Contigs with a Δbit-score lower or equal to -10 or with frameshift mutations were considered as putative EVE-like sequences and were further on excluded for segment assemblies. This cut-off was set to allow the exclusion of contigs with a higher amino-acid similarity to EVE sequences but would not exclude contigs with a similar low identity to both databases. This threshold was chosen empirically and should be taken cautiously.

### Contig extensions and post-processing

Nucleotide reference sequences resulting in a positive hit for orthomyxoviruses were downloaded from NCBI GenBank with efetch (v14.6). Contigs were mapped to a non-redundant referencing sequence list corresponding to their best blastx hits in Geneious Prime (v.2022.1.1). Contigs spanning only parts of the Orthomyxovirus segments were artificially extended to the full segment length by merging them with their reference sequence determined by their best blast hit. Non-host reads used for assembly were mapped back strictly (untrimmed) to the chimeric contigs using Bowtie2 (v2.3.5.1). The generated alignment file was converted, sorted and indexed using Samtools (v1.10, htslib 1.10.2-3) with a mapping quality filter of MAPQ>= 2. Coverage and sequencing depth were assessed using bedtools (v2.27.1). A sequence depth < 3 was considered insufficient for subsequent analyses and nucleotides were replaced with ambiguous Ns. Single nucleotide variants were called using Lofreq* (version 2.1.2) and sequences were corrected when variant abundance reached >50%. Coverage lengths and segment distribution per sample were illustrated with the ggplot2 function geom_tile (v3.3.5).

### Phylogenetic analyses

Potential viral proteins, EVEs and corresponding homologs of Orthomyxovirus reference sequences were aligned with the Geneious implemented MAFFT aligner (v7.490) employing either the L-INS-i, FFT-NS-i or FFT-NS-2 algorithm, which was automatically adjusted to the input query (Katoh & Standley, 2013). Because of the highly divergent nature of the sequence alignment, ambiguously aligned regions and gapped sites were pruned manually in Geneious. Sequences with more than 50% unknown/missing residues (represented as X in amino-acid sequence) were excluded from the final alignment but can be found in the Supplementary Data. Due to most sequences being partial (either through sequence depth or as an EVE) no strict alignment lengths were applied. Phylogenetic trees were constructed using IQTREE (version 1.6.12) (Nguyen et al., 2015). The substitution models were selected based on the Bayesian information criterion provided by the IQTREE-implemented ModelFinder (Kalyaanamoorthy et al., 2017): Segment PB1: LG+F+I+G4, Segment PB2: LG+F+G4, Segment PA: LG+F+I+G4, Segment NP: VT+I+G4, Segment GP: VT+I+G4, Segment HP1: VT+G4, Segment HP2: FLU+F+G4, Segment HP3: VT+G4. Branch support values were measured as ultrafast bootstrap by UFBoot2 with 1000 replicates (Hoang et al., 2018). The trees were visualized with the ggtree package (version 2.2.1) (Yu et al., 2018). Multiple alignments and Newick tree files generated by phylogenetic analyses are listed in Supplementary Data.

## Results

### Exploration of *Orthomyxovirus*-associated EVEs in genomic datasets

To explore the impact of EVEs on virus discovery, we conducted a targeted case study examining orthomyxoviruses in Culicinae mosquitoes. Our initial step involved determining the presence of EVEs within 13 publicly available genome assemblies containing the species *Culex pipiens* (n=1*), Culex quinquefasciatus* (n=2), *Aedes aegypti* (n=5), and *Aedes albopictus* (n=5), with some of them corresponding to strains and isolates widely used in cell culture (Supplementary Table 1). The screening identified *Orthomyxovirus*-like sequences in 10 of the 13 genomes, all corresponding to the *Aedes* genus. We identified a total of 223 significant matches (e-values <1×10^-4^) to orthomyxoviruses in the genomes of *Aedes aegypti* and *Aedes albopictus* (Supplementary Table 3). No matches were found within *Culex* genomes. EVEs related to 17 species were identified, all assigned to unclassified species within the genus *Quaranjavirus*. Most detected EVEs reveal numerous genomic sequences related primarily to the Guadeloupe mosquito quaranja-like virus, Usinis virus, and Aedes alboannulatus orthomyxo-like virus. The reported sequences have 26–91% (average 51%) amino acid (aa) identity with the present-day known exogenous *Orthomyxovirus* proteins. 81% of EVEs were derived from the nucleoprotein (NP) gene (n=180), while others derived from the RNA-dependent-RNA-polymerase (RdRP) associated segment PB1 (n=6), the glycoprotein (n=9) and hypothetical proteins 2 (n=23) and 3 (n=5) with currently unknown function (Figure 2). 12 EVEs related to the nucleoprotein, glycoprotein and hypothetical protein 3 encompass the entire length of the segment. Overall coverage lengths of EVEs compared to the corresponding reference sequence vary from 11.3%-100%, representing partial and full segment integrations (Supplementary Table 4).

**Figure 2.**
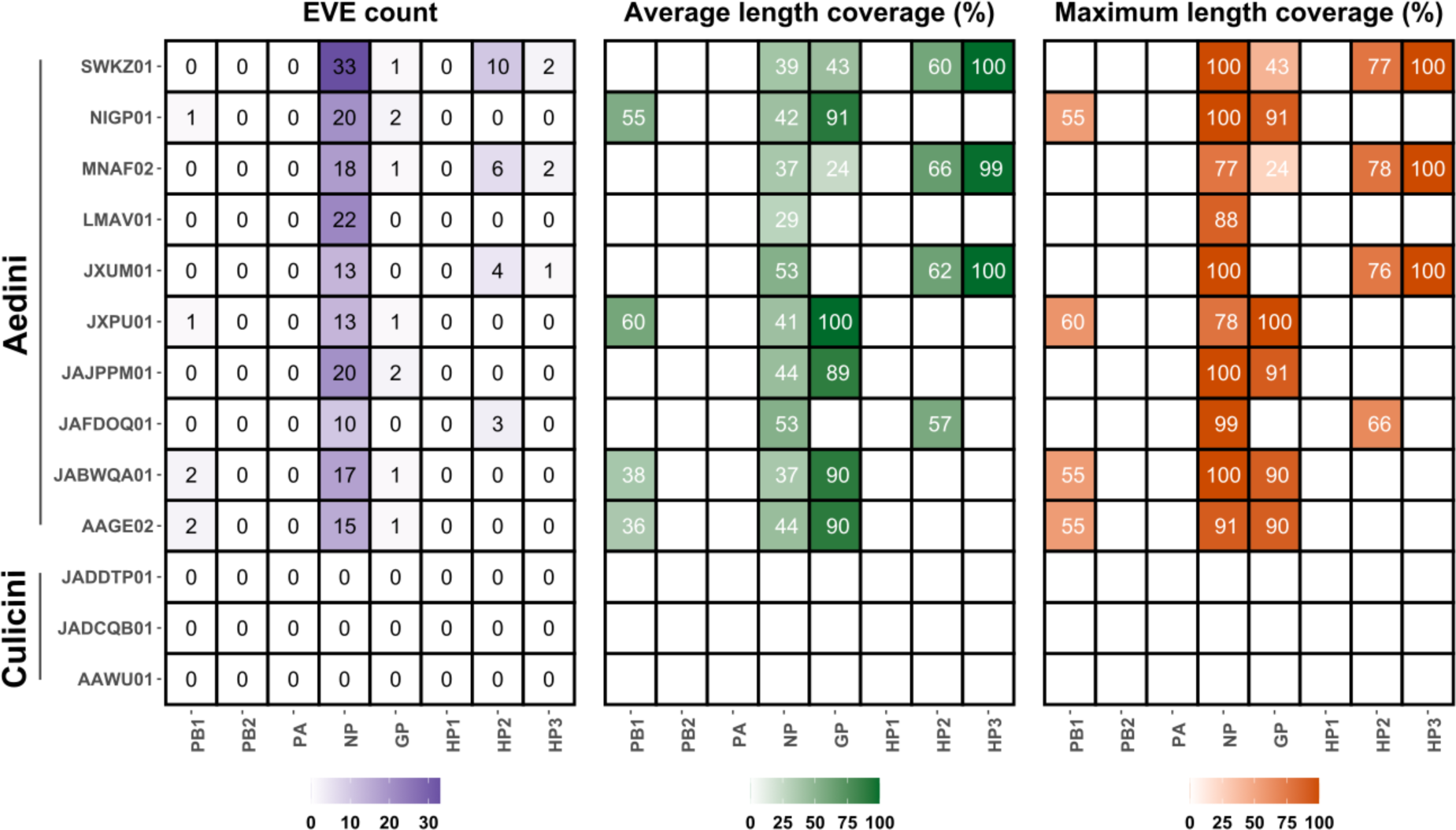
EVEs within genomic datasets exhibit bias toward nucleoprotein integration and showcase full ORF coverages. Tiles of heatmaps represent Orthomyxovirus segments of 10 Aedini and 3 Culicini mosquito genome assemblies (top to bottom). Length coverages were calculated by dividing EVE hits’ lengths (in amino acids) by the length (in amino acids) of the closest Orthomyxovirus reference hit. Detailed descriptions of mosquito datasets and EVE length coverages can be found in Supplementary table 1 and 4, respectively.

In the predicted coding sequence, 122 sequences contained at least one frameshift mutation and 85 EVEs contained premature stop codons. However, 73 EVE hits (32.3% of all genomic EVEs discovered) did not contain stop codons or frameshifts and can be considered intact full or partial ORFs (Supplementary Table 4). In addition, six hits were roughly the same size (>90% amino-acid length) as the related viral segments.

In addition, the reference genomes and transcriptomes, later used for host read removal in transcriptomics dataset analyses, were examined for EVEs. *Culex* datasets showed no evidence of EVE sequences. However, in the *Aedes* datasets, 33 EVEs were identified in the genome, and the transcriptome contained 4 EVEs identical to the genomic EVEs (Supplementary table 5). These EVEs exhibited amino acid identities ranging from 24% to 82% to exogenous *Orthomyxovirus* proteins. Among the identified EVEs, 84% were derived from the nucleoprotein (NP) gene (n=28), while the remaining EVEs originated from the RdRP-associated segment PB1 (n=2), hypothetical protein 2 (n=2), and the glycoprotein (n=1).

### Screening for orthomyxoviruses in transcriptomic datasets

To understand how EVEs may affect virus detection in transcriptomic data, we sought to identify transcribed EVEs and external viral sequences associated with Orthomyxoviridae in publicly available RNA-Seq datasets. A total of 538 RNA-Seq libraries from 23 BioProjects were downloaded from the Sequence Read Archive (SRA) NCBI database (Supplementary Table 6) (Leinonen, Sugawara, and Shumway 2011). Approximately half of the BioProjects (n=12) have been previously used for virus detection (Batovska et al., 2019; Batson et al., 2021; Chandler et al., 2014; McBride et al., 2014; C. Shi et al., 2019), with three known to have annotated Orthomyxovirus accession entries (Supplementary table 7), but were screened nevertheless and considered controls for our virus discovery pipeline. Screened samples mainly represent wild-caught adults or larvae from individuals or pools from various geographic locations (Figure 3a). Tribes included in the transcriptomic data are represented by 166 Aedini, 321 Culicini, 2 Culisetini, 5 Mansoniini, 3 Sabethini, 6 Toxorhynchitini, 1 Uranotaeniini and 10 non-defined metatranscriptomic samples.

**Figure 3:**
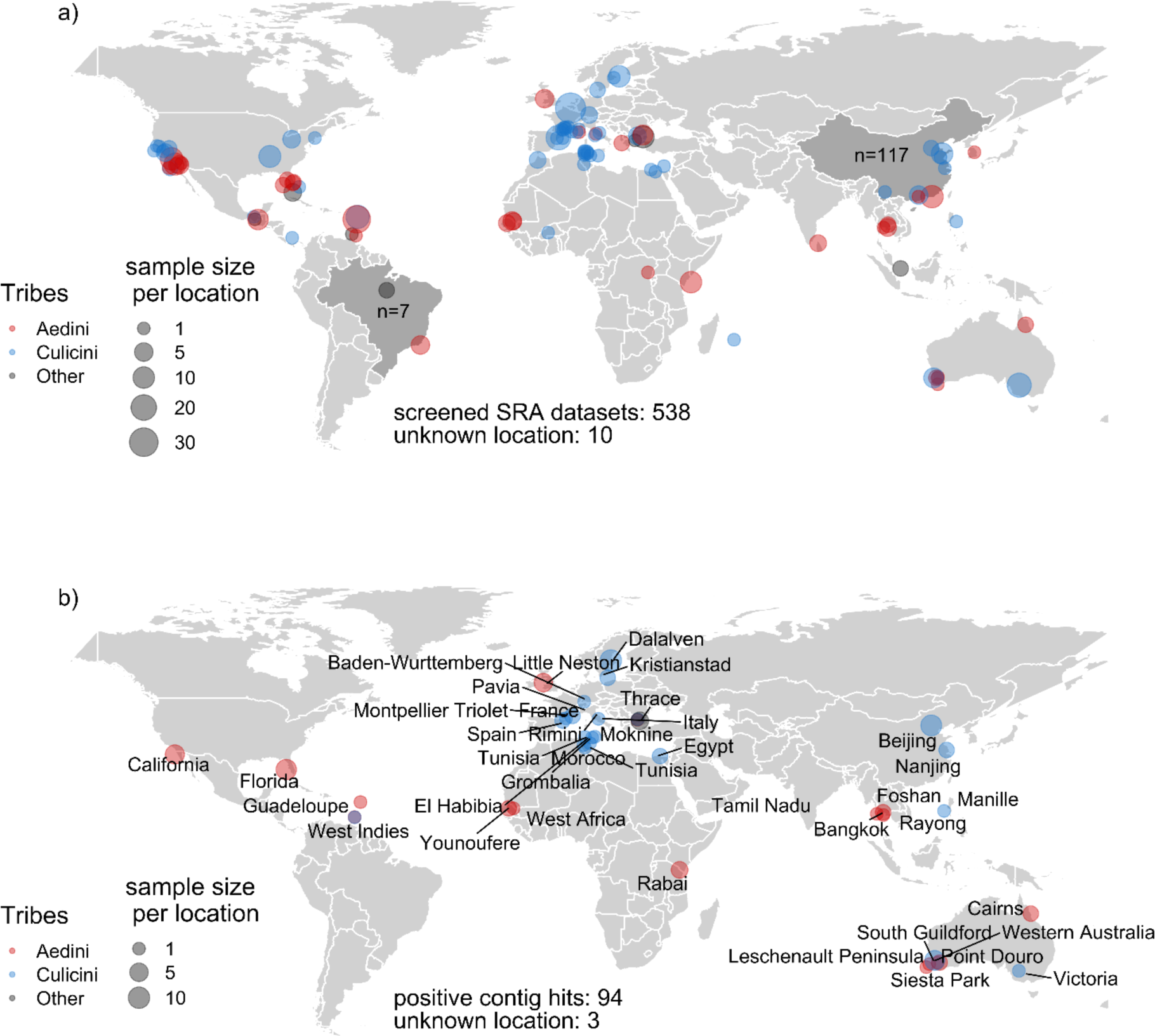
Positive samples of Orthomyxoviruses demonstrate a global distribution pattern. Red, blue and grey dots indicate the sampling site, provided by the original study, of *Aedes*, *Culex* and other Culicinae transcriptomic datasets screened. Size represents sample count. Latitude and longitude positions were either taken from metadata or estimated according to the location name. a) Screened datasets used for virus discovery pipeline. Dark grey areas (Brazil, China) indicate additional screened samples with unknown coordinates. b) Samples with positive *Orthomyxovirus* contig hits.

Orthomyxovirus-like contigs were present in 94 (∼18% of total datasets screened) samples, for which 72 samples have unannotated accessions for these viruses. These sequence hits were found in datasets belonging to the Culicinae tribes Aedini (42), Culicini (48) and 4 non-defined metatranscriptomic samples (Figure 3b). *Culex* samples are primarily located in Europe. *Aedes* samples (mostly tropical species) are centered around the tropics and sub-tropics. As expected, previously annotated *Orthomyxovirus* sequences in control datasets have been detected again by our pipeline.

The screening of sequencing libraries resulting from host read removal without integrated EVEs identified 2,494 contigs with *Orthomyxovirus*-like hits (Supplementary tables 8 and 9). The majority of the contigs were taxonomically assigned to yet unclassified species of the genus *Quaranjavirus* detected primarily in mosquito hosts. Guadeloupe mosquito quaranja-like virus 1 and Wuhan Mosquito Virus 6 were the most represented, with over 600 hits each. We identified 25 samples (without controls) where we could detect all 8 segments. However, additional *Orthomyxovirus* contigs with distinct nucleotide similarities to the full-length segments were detected for segments in the majority of the samples (Figure 4 and Supplementary Table 10). 7 Aedini samples comprised only sequences associated with the nucleoprotein (Figure 4), and 34 samples contained additional sequences per segment. 19 datasets associated with *Culex* mosquitoes contained only 3 segments or less. These sequences mostly showed low similarity to their best reference hit, incomplete fragment lengths and often mapped to the same loci (data not shown). Besides NP, these were primarily associated with the RdRP catalytic subunit 1 (PB1), the glycoprotein (GP) and the hypothetical proteins HP1 and HP3.

**Figure 4:**
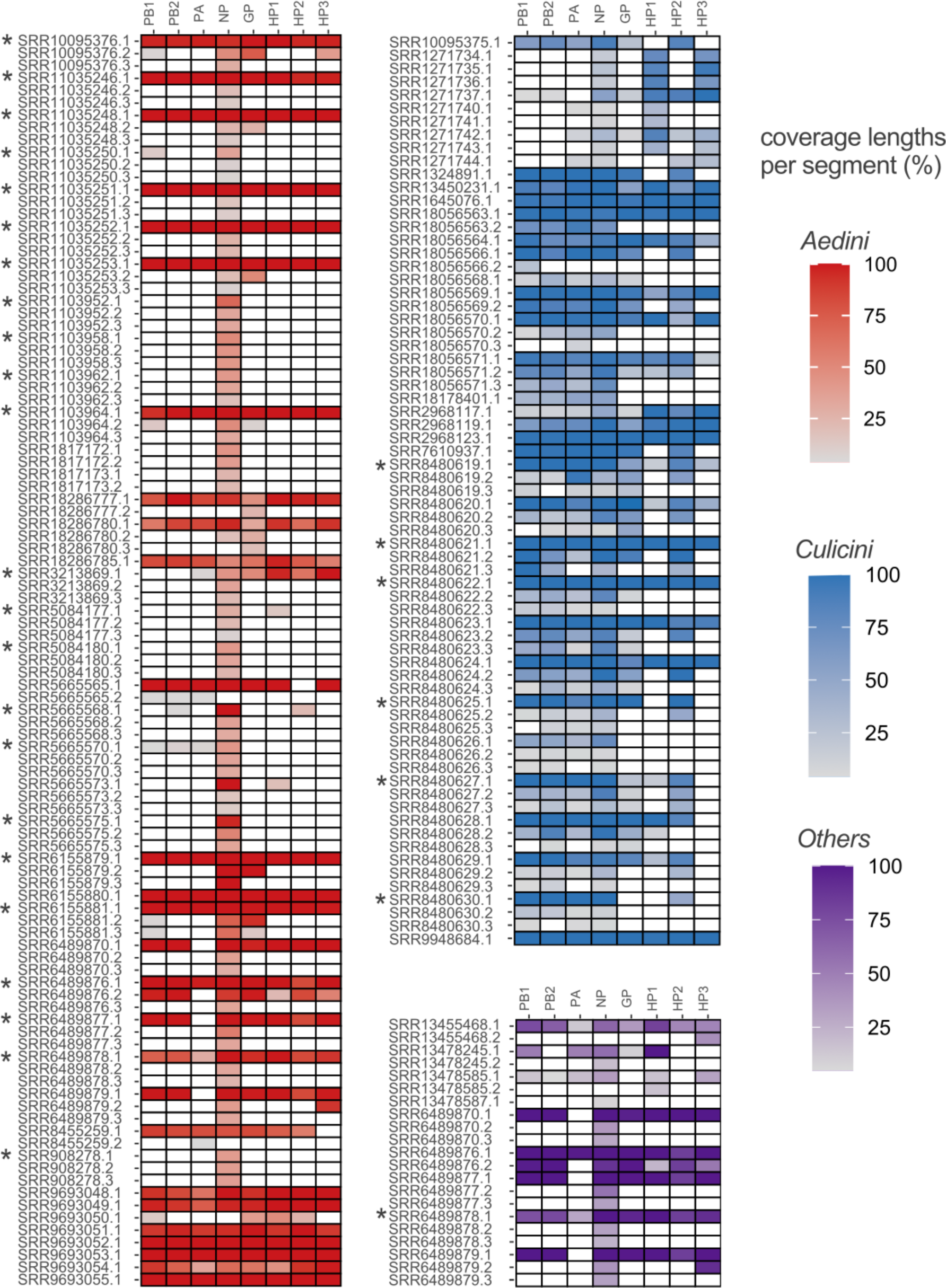
*Orthomyxovirus* detection in SRA datasets reveals an abundance of isolated segments. Here, we assess the completeness of our contigs for each viral segment. Contigs are aligned to each segment’s best matching accession hits to check for full or partial segment lengths. Additional contigs with high sequence variety to an already aligned sequence per segment were separated and treated as additional segments. For each segment length coverages were calculated. Aedini, Culicini and other Culicinae samples are colored in a red, blue, or violet gradient according to their coverage percentages, respectively. Segments per sample were ordered according to their highest percentages and are represented as tiles. Appended numbers to accession numbers (.2, .3) represent additional sequences per segment per sample, with up to three additional sequences shown in this figure. Samples with more than three sequences per segment are annotated with an asterisk and complete coverage lengths for sequences can be found in Supplementary Table 10. Segments with no associated contigs are depicted in white.

### EVE detection in transcriptomic data

To explore the potential presence of EVE-like sequences within transcriptomic data analysis, we generated two custom databases consisting of either the already existing *Orthomyxovirus* reference list or the translated amino-acid sequences of previously detected genomic EVEs. Since genomic EVEs were only found in *Aedes* assemblies, only contigs with *Orthomyxovirus* hits derived from Aedini samples were used as a query for a diamond blastx search against the two databases. The highest bit score hits against each database were merged into a table and the difference between bit scores (Δ bit score) was calculated (Supplementary Table 11). The highest bit score values and Δ bit scores were plotted in Figure 5A/B. High overall bit scores for non-EVE (exogenous Orthomyxovirus references, maximum bit scores >500) sequences also display a higher Δ bit score (positive values). This indicates that putative exogenous sequences have hits with longer length matches to non-EVE references, shorter length matches to EVE sequences, and significantly lower amino-acid similarities to EVEs. These long-length matches are, with a few exceptions, missing for hits with a better bit score for genomic EVE sequences.

**Figure 5:**
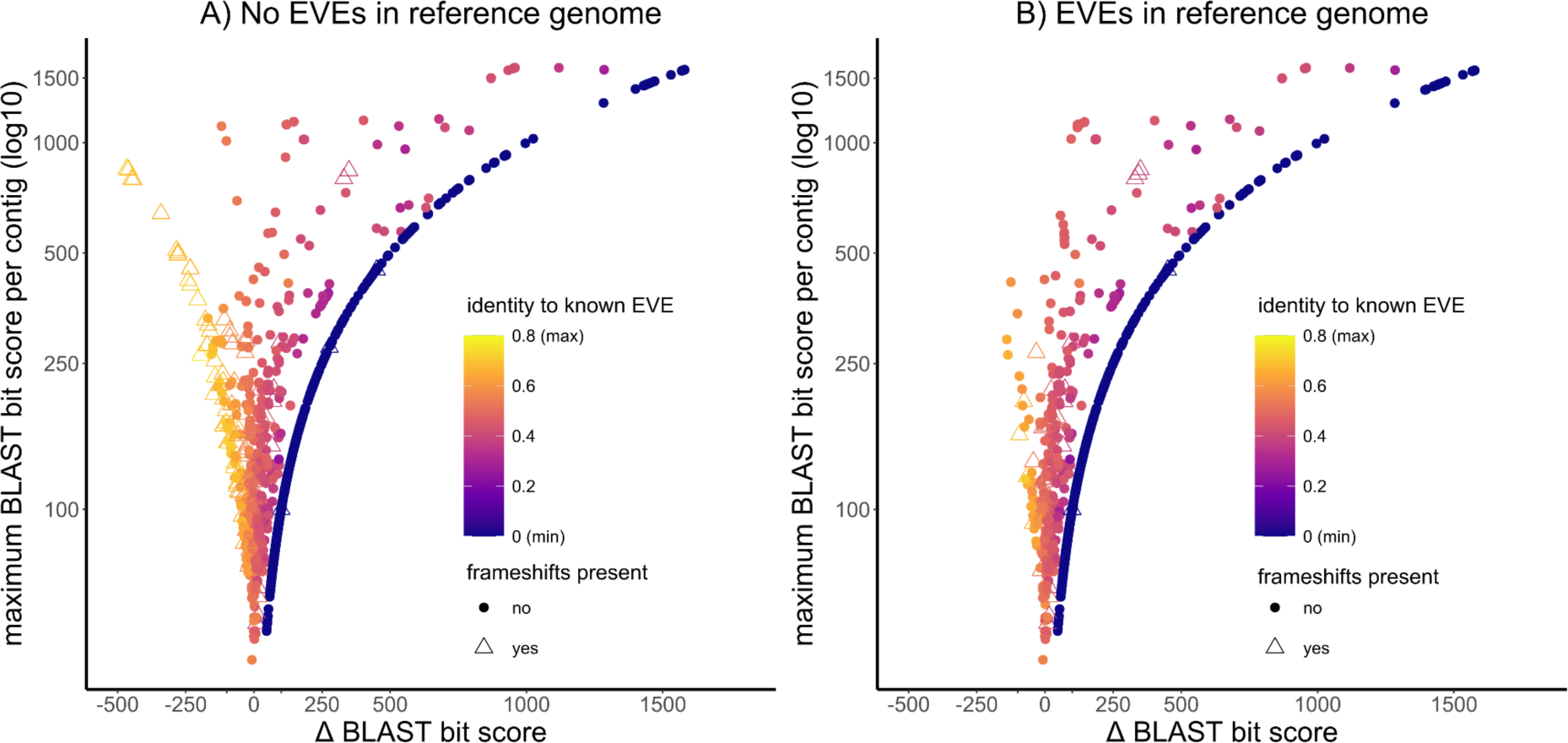
Bit score comparisons of SRR contig hits against EVE or exogenous viral sequences show reduced hits for pipeline with host read removal step, including EVE sequences. Contigs were blasted against two custom databases: the existing *Orthomyxovirus* reference list or the translated amino-acid sequences of previously detected genomic EVEs. Data points are colored according to amino-acid sequence similarity to EVE references. Contigs with frameshifts or nonsense mutations (stop codons) are indicated with triangular shapes. Contigs after host read removal without integrated EVEs in the reference genome are shown in panel A. Contigs after host read removal with present EVEs in the reference genome are shown in panel B. Details of the bit score calculations and frameshift corrections can be found in Supplementary Table 11.

Since most of the screened datasets were not generated for virus detection, the quality of viral sequence reads is generally low, which leads to shorter read assemblies. Consequently, maximum bit scores are low and show little difference in amino-acid identity when aligned to EVEs or non-EVEs (-100 < Δ bit score < 100). As different library preparation techniques were used for different BioProjects, bit score data points were analyzed by BioProject (Supplementary Figure 1), but no correlations between Δ bit score distributions and virus- or non-virus-focused studies could be observed. Figure 5 also depicts detected frameshifts or nonsense mutations in *Aedes*-associated contigs. While most of such mutations occur at negative Δ bit scores (more EVE-related), some contigs with higher similarity to exogenous reference sequences also exhibit frameshifts. As bit score calculations are dependent on identified genomic EVEs, false positives for exogenous-like sequences are to be expected.

### Influence of EVEs on host read removal

Upon host read removal from reference sequences with EVEs, 213 viral contigs were lost compared to contigs generated from reads passing host read filtering without EVEs. As shown in Figure 5B, most of the lost contigs (n=107) have higher blast bit scores for genomic EVEs (Δ bit score lower/equal -10) than exogenous *Orthomyxovirus* sequences. However, 45 contigs associated with the exogenous virus (Δ bit score greater than -10 + no frameshift/stop codons) were also lost. Additionally, not all EVE-associated sequences were lost due to host read removal, as contigs with frameshift/nonsense mutations can still be seen within the plot (Figure 5B).

To avoid chimeric assemblies between putative EVE and non-EVE sequences during the post-processing analysis, a Δ bit score cut-off was used. Contigs with a Δ bit score lower than -10 and frameshift mutations (n=179) were set aside as potential EVE sequences and excluded from manual contig extensions. Out of these, 84 contained frameshift mutations, and 12 contained nonsense mutations (stop codons) (Supplementary table 11).

### Influence of EVEs on read coverage and SNP analysis

For a more profound understanding of contig loss during host read removal, we closely inspect this process at the read level, conducting read coverage analysis across all four workflows. After mapping on the host genome containing EVEs, a reduction of read abundance can be seen in 22 NP segments (68% of *Aedes* samples). Upon comparison of host read removal with the different transcriptomic references, no changes in sequencing depth were observed. We compared nucleotide similarities between genomic EVEs and our transcriptomic datasets to investigate further the potential bias of EVE presence in host read removal approaches. Regions with reduced read depth were isolated and aligned against EVE sequences found in the reference genome. Best-aligned hits can be seen in Supplementary Table 12. Nucleotide similarities ranged between 74%-100%. To elucidate the impact of read reduction on the overall loss of sequence information, coverage comparisons between the four generated sequences from each workflow were done and can be found in Supplementary Figure 2. Three sequences exhibited regions within the segments that fell below the sequencing depth threshold after genomic host read removal with EVE presence. Lastly, single nucleotide polymorphisms (SNPs) were explored among the four different pipelines. While SNPs could be detected in 20 datasets, most can be attributed to equally abundant variants chosen stochastically. However, 4 SNPs generated within regions of reduced read depth show a switch between major/minor variants and are therefore attributed to differences in host mapping (SNP distributions in Supplementary Figure 2).

### Phylogeny and implications on taxonomy

To illustrate the evolutionary relationships between exogenous orthomyxoviruses and EVEs, we aligned putative EVE protein sequences with viral protein sequences obtained after removing host reads containing EVEs. We then constructed maximum likelihood phylogenetic trees for each genetic segment (Supplementary Figures 3-9, Figure 6). The nucleoprotein was chosen as representative, as most SRA samples and EVEs correspond to this segment (Figure 6). The final sequences produced in this study clustered in two distinct clades, forming a sister lineage to the classified genus *Quaranjavirus*. The first clade (top) consists of primarily *Aedes*-associated orthomyxoviruses except for Astopletus virus, a recently identified Quaranjavirus detected in Culicini. 18 transcriptomic samples clustered with Guadeloupe mosquito quaranja-like virus 1, detected initially in Guadeloupe and California (Batson et al., 2021; C. Shi et al., 2019). 6 samples clustered to Aedes detritus orthomyxo-like virus and one to Aedes alboannulatus orthomyxo-like virus. For the second clade (bottom), 19 datasets include sequences identified as Wuhan Mosquito virus 6 and Culex pipiens orthomyxo-like virus. While these two references have been deposited with different names and are derived from different continents, they share a >99% amino-acid similarity. Four datasets from the same BioProject clustered on a separate branch to the known Wuhan mosquito viruses 6. 16 sequences were designated as Wuhan Mosquito virus 4-like and 16 sequences clustered with Culex orthomyxo-like virus.

**Figure 6:**
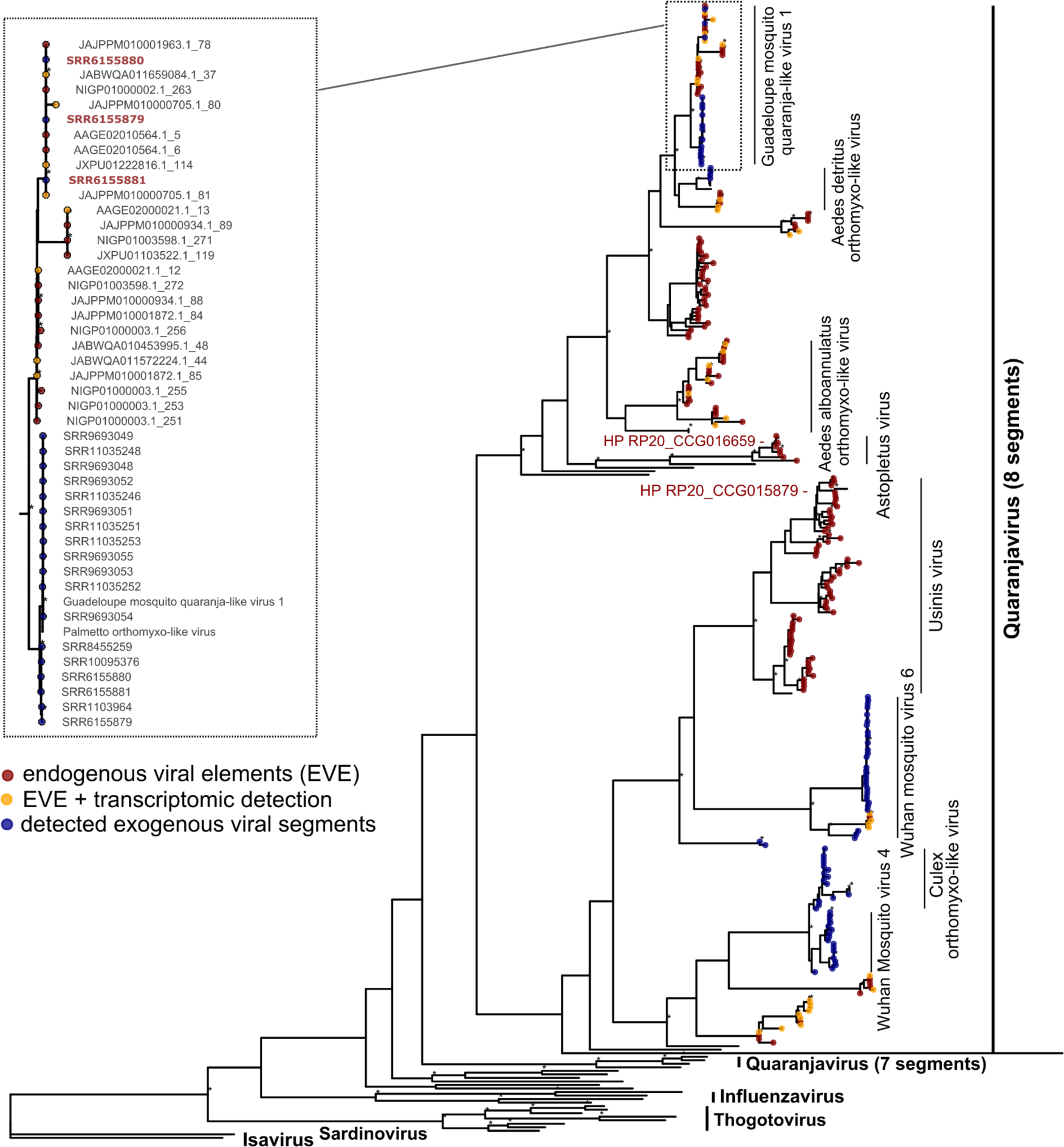
Phylogenetic tree depicting relationships between NP sequences of orthomyxoviruses. Sequence alignments of genomic EVEs, transcriptomic orthomyxovirus sequences, and references were generated with mafft v7.490. The maximum likelihood tree was constructed with model VT+I+G4 in IQ-TREE 1.6.12 (+ ultrafast bootstrap (1000 replicates)). Significant ultrafast bootstrap values >95% are depicted with an asterisk. Protein sequences detected in this study are marked as colored circles at the tips: red - genomic EVEs, orange - genomic EVEs + detection of transcriptomic contigs with sequence similarity >95% and a Δbit score <-10, dark blue - putative exogenous sequences. Species and genus (bold) characteristics per clade are highlighted with black vertical lines to the right of the tree. Transcribed EVE sequences with fully intact ORF are depicted in bold red. Two hypothetical proteins from *Aedes albopictus* (HP RP20) are pointed out in red.

Genomic EVEs cluster within distinct sister clades to their exogenous viral counterparts and are closely related to each other. The 29 EVEs form three subgroups related foremost to the Guadeloupe mosquito quaranja-like virus 1. Six integrations show the highest similarity to the Aedes detritus orthomyxo-like virus, while 26 EVEs form a sister clade to the common ancestor of the two previously mentioned viral species. 25 EVEs cluster with Aedes alboannulatus orthomyxo-like virus, and 7 EVEs cluster with the recently described Astopletus virus. With 57 EVEs detected for Usinis virus, these sequences formed the most extensive clade of EVEs within the tree. Surprisingly, no new potential exogenous samples and putative transcriptomic EVEs could be found for the Usinis virus. 5 and 19 EVEs were closest related to Wuhan Mosquito virus 6 and 4, respectively. Astopletus virus, Wuhan Mosquito virus 4 and 6 have only been detected in Culex mosquitoes previously. However, all detected genomic EVEs were detected in Aedes genomic data, which suggests that they are or were able also to infect Aedes mosquitoes persistently.

In addition to genomic EVEs, putative transcribed endogenous viral contigs, previously excluded for further downstream analysis, with a high sequence similarity to genomic EVEs, were also visualized in the tree (depicted in orange). The excluded contigs (n= 209), had the highest similarities to 42 EVEs, all derived from *Aedes aegypti* host genomes, and can be found evenly distributed among the genomic EVE clades (except Usinis virus). To illustrate a lack of putative EVE filtering, three consensus sequences generated of putative EVE contigs (Δ bit score < -10) were included for subsequent sequence correction, coverage analysis, and tree construction. All consensus sequences depicted full-length contigs for an additional NP segment to an already complete *Orthomyxovirus* genome (all eight segments present) (Supplementary Figure 10), but also included extended sequences with blast hits to uncharacterized insect proteins, validating them as transcribed EVEs. As seen in Figure 6, the sequences, named after their SRA datasets SRR6155879, SRR6155880 and SRR6155881 (bold red), cluster within the genomic EVEs related to Guadeloupe mosquito quaranja-like virus 1. Upon closer inspection of their amino acid sequence, all sequences share 100% similarity to a detected genomic EVE with >98% segment coverage and a mean depth of 10, 22.6 and 36.2. All three sequences represent a fully intact ORF, share 100% identity, and, although derived from the same BioProject, differ in sampling locations (Cairns, Australia and Bangkok, Thailand), indicating a highly conserved integration event across geographically diverse samples.

While screening for genomic and transcriptomic integration events, various orthomyxoviral sequences had the highest or second-highest blast hits to hypothetical proteins annotated as part of the *Aedes albopictus* genome. This is true for EVEs related to Astopletus and Usinis viruses and both recently described Quaranja-like viruses with 8 segments (Batson et al., 2021). These proteins (NCBI accession numbers: KXJ73423.1 and KXJ73043.1) are likely Orthomyxovirus EVEs, falsely annotated as part of the host genome (Figure 6 and Supplementary Figure 11).

## Discussion

### EVEs and virus discovery

Viral metagenomic analysis has become an important approach to uncovering novel viral genomes in various samples, giving an ever-increasing insight into the diversity of the virosphere. With the rapid pace of improved sequencing techniques and assembly tools, viral sequence detection in samples has become sufficient evidence for admission as *bona fide* viruses, no longer relying on the verification through phenotypic properties.

While metagenomics and metatranscriptomics are undoubtedly the present and future of virus discovery, appropriate checks for data accuracy and integrity are essential to infer the actual existence of a virus with solely sequence data. Unfortunately, these steps are not standardized among virus discovery pipelines, leading to discrepancies regarding coverage and sequencing depth thresholds, ensuring the validity of viral sequences. In addition, more and more studies focus on re-analyzing samples initially intended for non-virus-related applications. Before and during library preparation, suboptimal processing often correlates with poor viral RNA quality, leading to lower read abundance and shorter assembled contigs. Complete genome assemblies are difficult, and scaffold assemblies and contig extensions are often necessary to reach full genome length. These assembled sequences risk being derived from multiple virus populations, leading to artificially generated chimeric genomes, which can occur between closely related exogenous virus populations and with integrated viral sequences in the host if their sequences are sufficiently similar.

Non-retroviral EVEs in the sequenced host genome are often ignored in virus discovery pipelines, and bioinformatic steps undertaking EVE removal are rare. RNA-Seq libraries especially forgo this additional downstream cleaning step, as DNA removal steps and mRNA enrichment techniques are often part of cDNA library preparations. However, some EVEs are transcribed and present among the sequenced transcribed RNAs (Katzourakis & Gifford, 2010). Our study shows that an important number of RNA sequences identified from RNA-Seq libraries have high amino acid similarities with EVEs detected from corresponding genomic host datasets. While most of these sequences only consist of partial segments or contain frameshift mutations, some of them can span whole segments and display intact open reading frames and sufficient abundance levels. Without initial EVE removal steps, these sequences can be easily misinterpreted as belonging to exogenous orthomyxoviruses, as seen in this study. Samples with low-quality viral contigs are challenging to classify as either more EVE- or exogenous-like, especially considering that some genomic EVEs were identical to their respective exogenous counterparts, with intact full-length ORFS, and/or expressed. Without a host genome and preliminary EVE characterization, and in a context of low viral reads abundance, which is common in metagenomic studies, this can lead to the misidentification of an EVE as an exogenous virus, especially when sequence assemblies contain incomplete genomes of novel viruses. We thus suggest that partial genomes or a limited number of viral segments should be viewed cautiously to avoid a potential misidentification in sequence origin.

When a host genome is available, reads are typically mapped to a host genome or transcriptome reference to alleviate the time-consuming assembly tasks. While this limits the presence of EVEs in the assembled viral contigs, it can also lead to loss of information as virus-associated reads can match genomic EVEs, as we have shown in this study. This can be especially detrimental for sup-optimal samples without virus enrichment, as read depths are usually low and could get lost during quality cut-offs. Despite not being a common event, integrated EVEs in the reference genome might also alter the final consensus sequences of the present viral population. This further complicates the already challenging task of distinguishing mixed infections or viral population clouds as potential variations at single nucleotide positions (SNPs) could be observed in our data.

### Orthomyxoviridae-derived EVEs in mosquitoes

Despite the limited sampling size of non-retroviral EVEs, most seem to arise from negative-strand RNA viruses (Aiewsakun & Katzourakis, 2015; Blair et al., 2020; Holmes, 2011). The exact mechanism of this favoritism of ssRNA− viruses is unclear, but one contributing factor could be the replication of certain viral families, such as Bornaviridae and Orthomyxoviridae, within the nucleus (Horie et al. 2010). This replication location might increase the likelihood of reverse transcription and subsequent genomic integration. However, only a limited number of *Orthomyxovirus*-derived EVEs have been described so far (Katzourakis & Gifford, 2010; Li et al., 2015), maybe due to limited knowledge on the diversity of the family, until recently mostly restricted to Influenzavirus and Thogotovirus. Recent studies have unveiled that orthomyxoviruses are commonly represented in arthropods (Batson et al., 2021; M. Shi et al., 2016). By including a diverse range of newly described orthomyxovirus species as a reference input, our study now expands the observation of *Orthomyxovirus*-derived EVEs to Culicinae host genomes. Similarly to EVE-derived from other viral families, detected *Orthomyxovirus*-derived EVE were found in *Aedes* mosquito genomes. Despite the high prevalence of *Orthomyxovirus* sequences in transcriptomic *Culex* datasets, no EVEs were detected in *Culex* genomes. The reason for such low levels of viral integration is not known yet but is speculated to correlate to the presence of transposable elements (TEs) (Whitfield et al., 2017). However, while *C. quinquefasciatus* genome is known to comprise only 29% of transposable elements (TEs) compared to *Aedes aegypti* (42-47%), the *Anopheles gambiae* genome has even fewer TEs (11-16%), but shows a higher number of EVEs (Arensburger et al., 2010; Blair et al., 2020; Holt et al., 2002; Nene et al., 2007).

As seen in other EVE studies, we observed preferential integration of viral genes or segments: sequences of EVEs encode for the viral RNA-dependent-RNA-polymerase subunit PB1, the nucleocapsid (NP), a glycoprotein and two hypothetical proteins. Considering that over 80% of the observed EVEs correspond to NP genes, a selective pressure to endogenize and/or keep this viral genomic region might be hypothesized. NP, the most abundant protein in infected cells (Kummer et al., 2014), has an essential role in viral RNA replication and transcription, organization of RNA packing, and nuclear trafficking (Eisfeld et al., 2015; Herz et al., 1981; Martin & Helenius, 1991). NP endogenization might thus result from relative mRNA abundance in the nucleus. Preferential integration of NP, but also polymerase genes, was also seen in other negative-sense RNA viruses (Gilbert & Belliardo, 2022; Katzourakis et al., 2007; Katzourakis & Gifford, 2010). However, since genome organization and replication mechanisms differ between virus families, the drivers of endogenization need to be explored further.

## Conclusions

Misidentification of sequences is expected to occur, considering the continuing exploration of viral sequencing data with metagenomics and metatranscriptomics. As seen in this study, annotation errors have already been detected in the genome of *Aedes albopictus*, with some hypothetical proteins being, in reality, integrated viral elements related to two recently discovered quaranjaviruses. In addition, misclassifying EVEs with exogenous viral sequences could contaminate databases and adversely affect downstream analyses, such as phylogenetic analyses.

To prevent this, some detection steps for EVEs in transcriptomic sequence data are possible. As we have shown, mapping reads on host genomes, when available, can alleviate this bias to some extent, albeit at the cost of sensitivity. Without a host genome, some ways to help distinguish EVEs from their respective exogenous counterparts include the detection of disrupted open reading frames, similarity to already known EVEs in genomic data in closely related species, transcript abundance compared to a housekeeping gene from the host, or whether the read contains any host DNA. Some computational tools like CheckV (Nayfach et al., 2021) can help filter some EVE sequences, but applications are limited due to sequence quality input requirements.

While there is no doubt that today’s virus discovery technologies heavily rely on metagenomic sequencing, it is essential to consider possible limitations due to the presence of EVEs in transcriptomic and meta-transcriptomic datasets. Datasets with a low viral abundance are especially susceptible to bias, primarily due to the challenge of distinguishing between exogenous and endogenous viral sequences. We argue that additional caution should be taken upon detecting novel virus sequences and that a framework for EVE detection should be a standard step within virus discovery pipelines and would provide great validation of exogenous viral sequences derived from metagenomic sequencing.

## Data availability statement

See https://github.com/nbrait/EVEs-bias-virus-discovery for a GitHub repository containing all scripts, intermediate files, sequences, tools and versions described in this paper. No raw sequencing data was generated for this study. Viral genomes are being submitted to NCBI GenBank; they can currently be accessed in the GitHub repository. All provided material will be submitted to a third-party repository upon manuscript publication.

## Author contributions

NB and SL conceived the study. NB performed formal analyses and contributed to visualization. NB, TH and SL contributed to validation. CM, AE and SG provided sequence input. SL contributed to supervision, project administration and funding acquisition. NB, TH and SL wrote the original draft of the manuscript. All authors read and approved the manuscript.

## Supplementary Figures

**Supplementary Figure 1:**
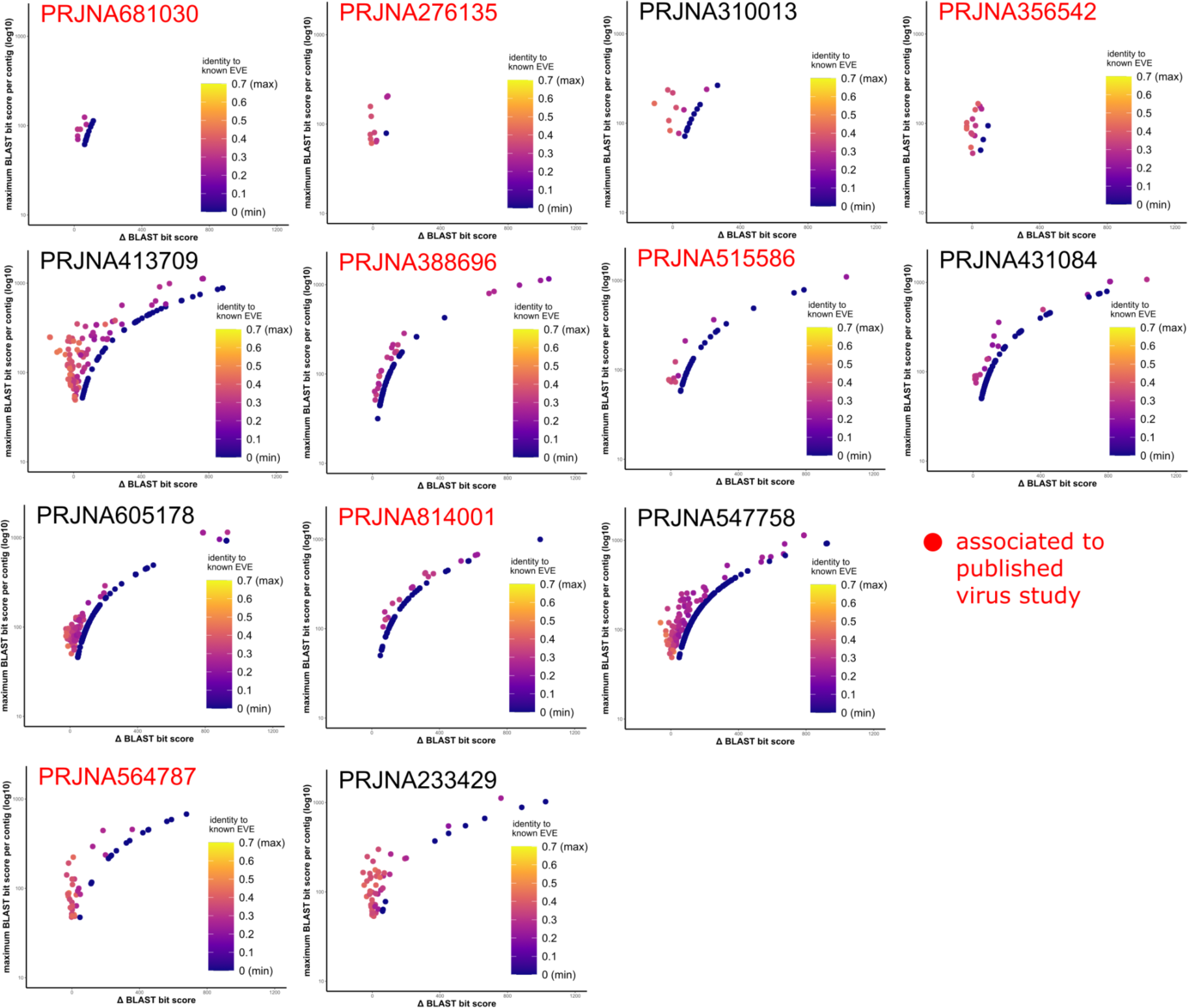
Plotted contig bit scores per bioproject. Contigs were blasted against two custom databases: the existing *Orthomyxovirus* reference list or the translated amino-acid sequences of previously detected genomic EVEs. Data points are coloured according to aa sequence similarity to EVE references. Bioprojects associated with published virus studies are indicated in red. R script and details to bit score calculations and frameshift corrections can be found in Supplementary table 11.

**Supplementary Figure 2.**
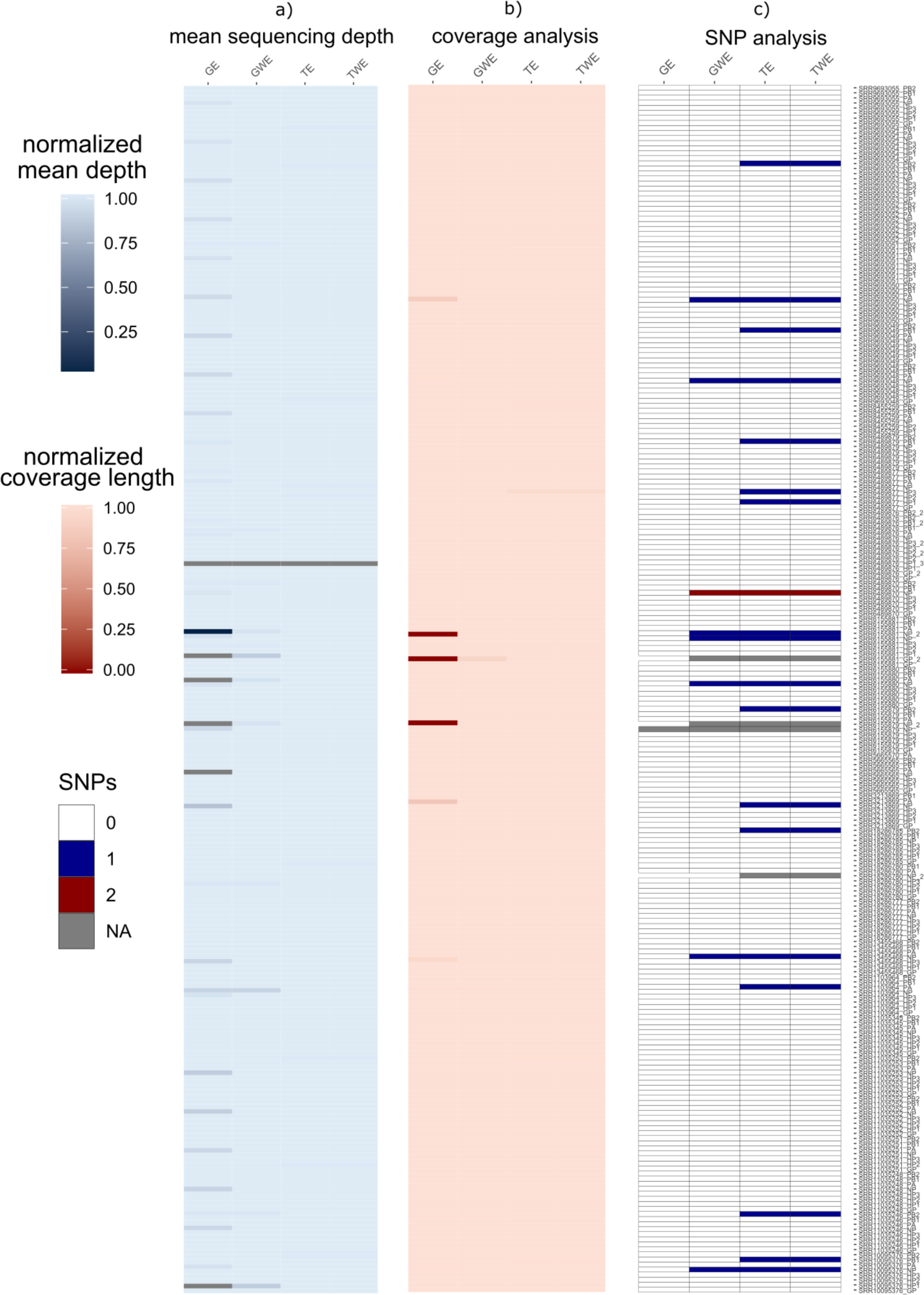
Comparison of mean sequence depths, coverage, and SNPs. Sequence reads from four different pipelines were mapped back to the chimeric reference sequences generated in Geneious Prime. Sequencing depths were generated with samtools. The percentages of length coverage were calculated by dividing the corrected coverage lengths minus the count of Ns by full coverage length. For SNP calculations, GE (genomic host read removal with EVE presence) was used as a reference sequence to observe differences in the other three workflows. a) Mean sequencing depth analysis, b) coverage analysis, c) SNP analysis. Abbreviations: GE (genomic host read removal with EVE presence), GWE (genomic host read removal without EVE presence), TE (transcriptomic host read removal with EVE presence), TWE (transcriptomic host read removal without EVE presence). Segments that did not generate sam output files after bowtie2 mapping are depicted in gray.

**Supplementary Figure 3:**
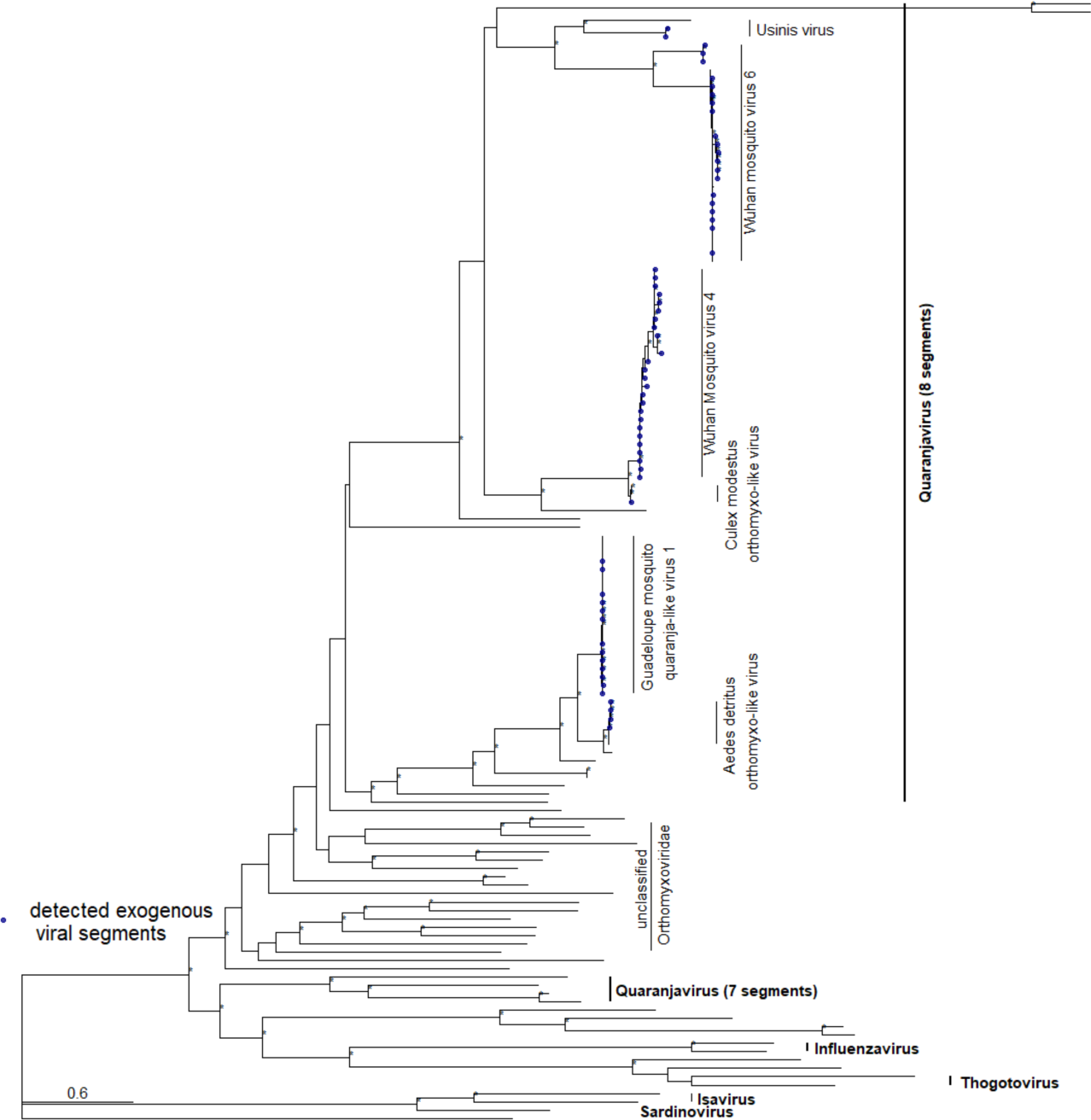
Phylogenetic tree depicting relationships between PB2 sequences of orthomyxoviruses. The maximum likelihood tree was constructed with model VT+I+G4 in IQ-TREE 1.6.12 (+ ultrafast bootstrap (1,000 replicates)). Significant ultrafast bootstrap values >95% are depicted with an asterisk. Detected protein sequences are marked in blue. Species and genus (bold) characteristics per clade are highlighted with black vertical lines to the right of the tree.

**Supplementary Figure 4:**
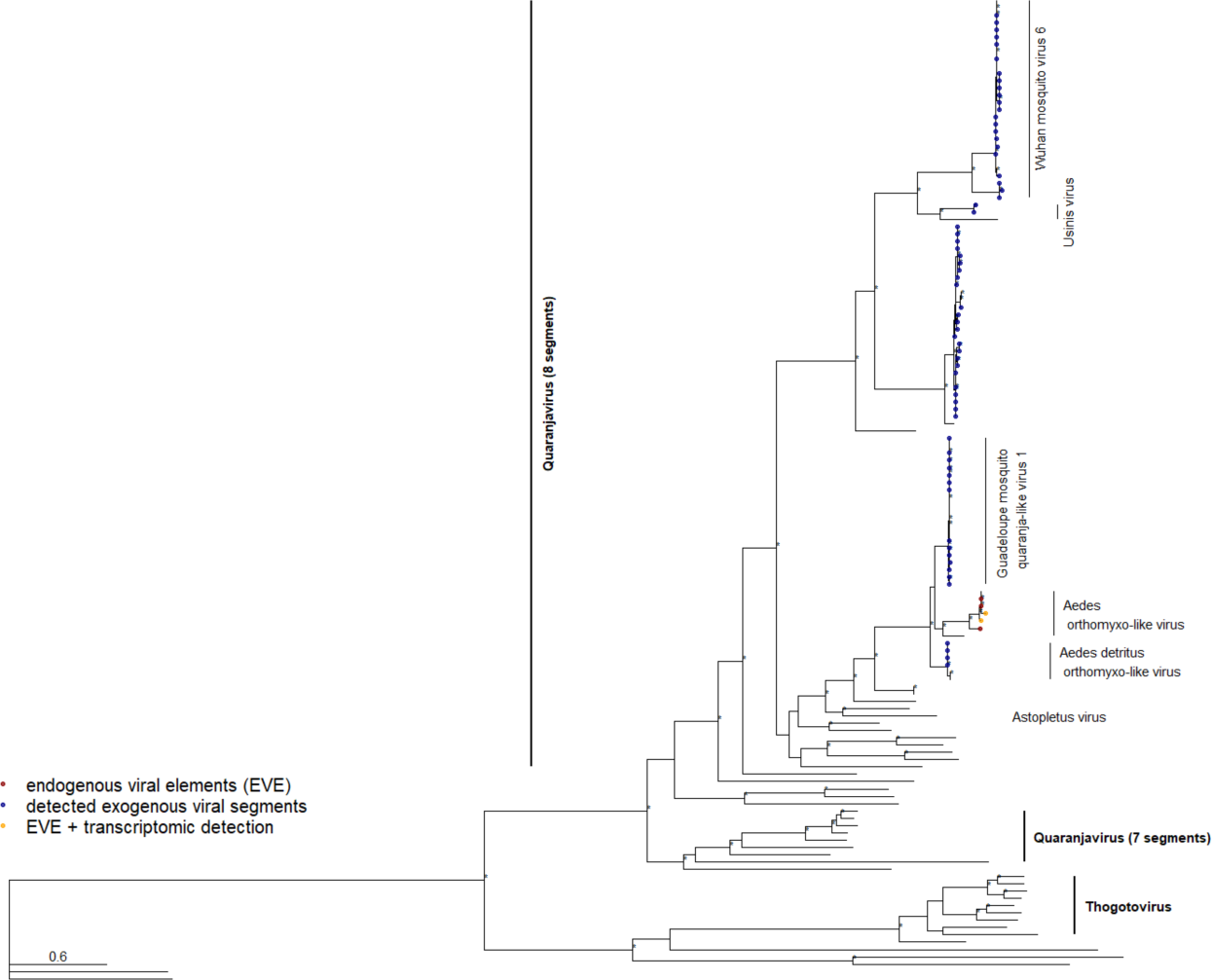
Phylogenetic tree depicting relationships between PB1 sequences of orthomyxoviruses. The maximum likelihood tree was constructed with model VT+I+G4 in IQ-TREE 1.6.12 (+ ultrafast bootstrap (1,000 replicates)). Significant ultrafast bootstrap values >95% are depicted with an asterisk. Detected protein sequences are marked in blue. Species and genus (bold) characteristics per clade are highlighted with black vertical lines to the right of the tree.

**Supplementary Figure 5:**
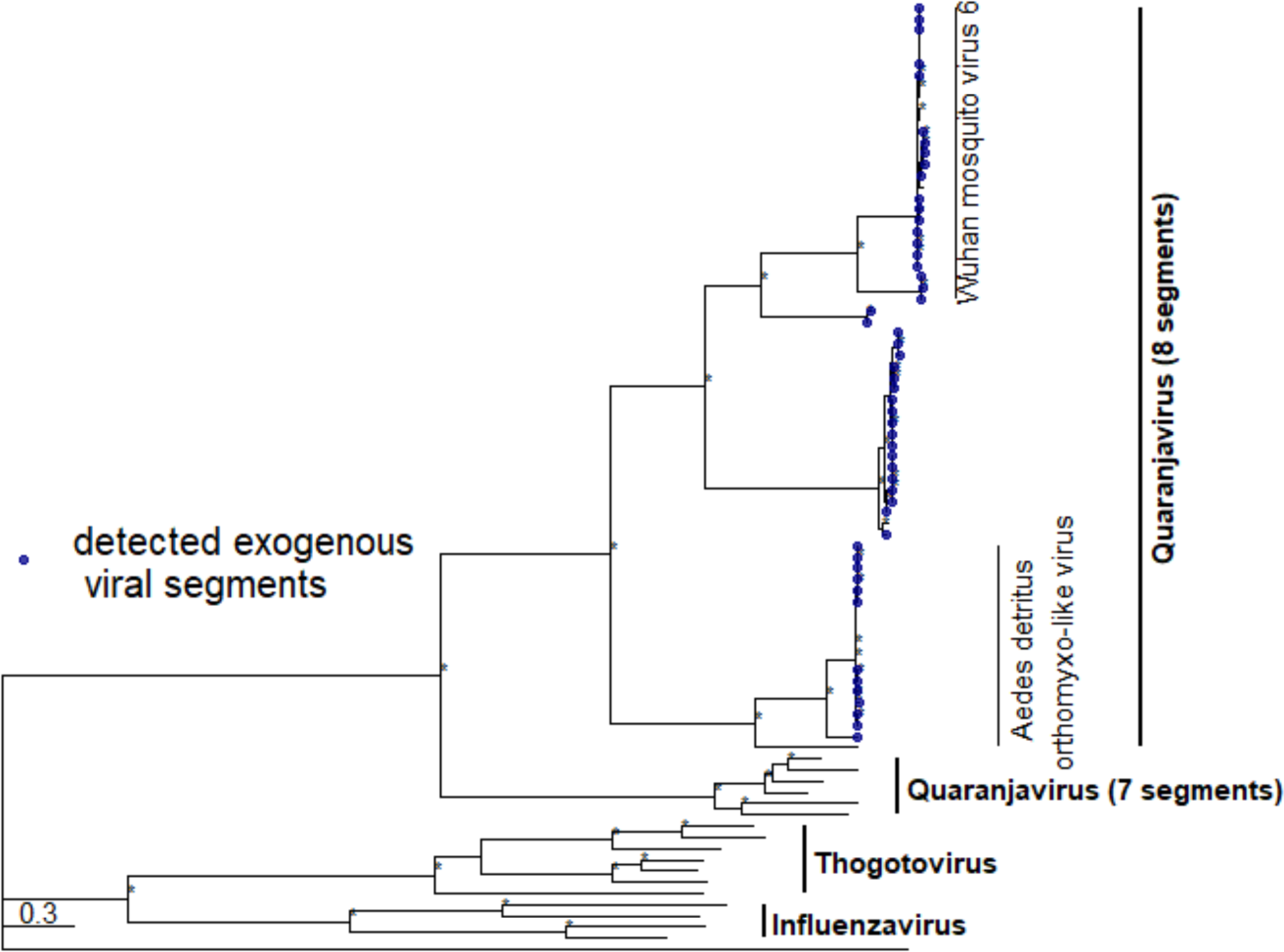
Phylogenetic tree depicting relationships between PA sequences of orthomyxoviruses. The maximum likelihood tree was constructed with model VT+I+G4 in IQ-TREE 1.6.12 (+ ultrafast bootstrap (1,000 replicates)). Significant ultrafast bootstrap values >95% are depicted with an asterisk. Detected protein sequences are marked in blue. Species and genus (bold) characteristics per clade are highlighted with black vertical lines to the right of the tree.

**Supplementary Figure 6:**
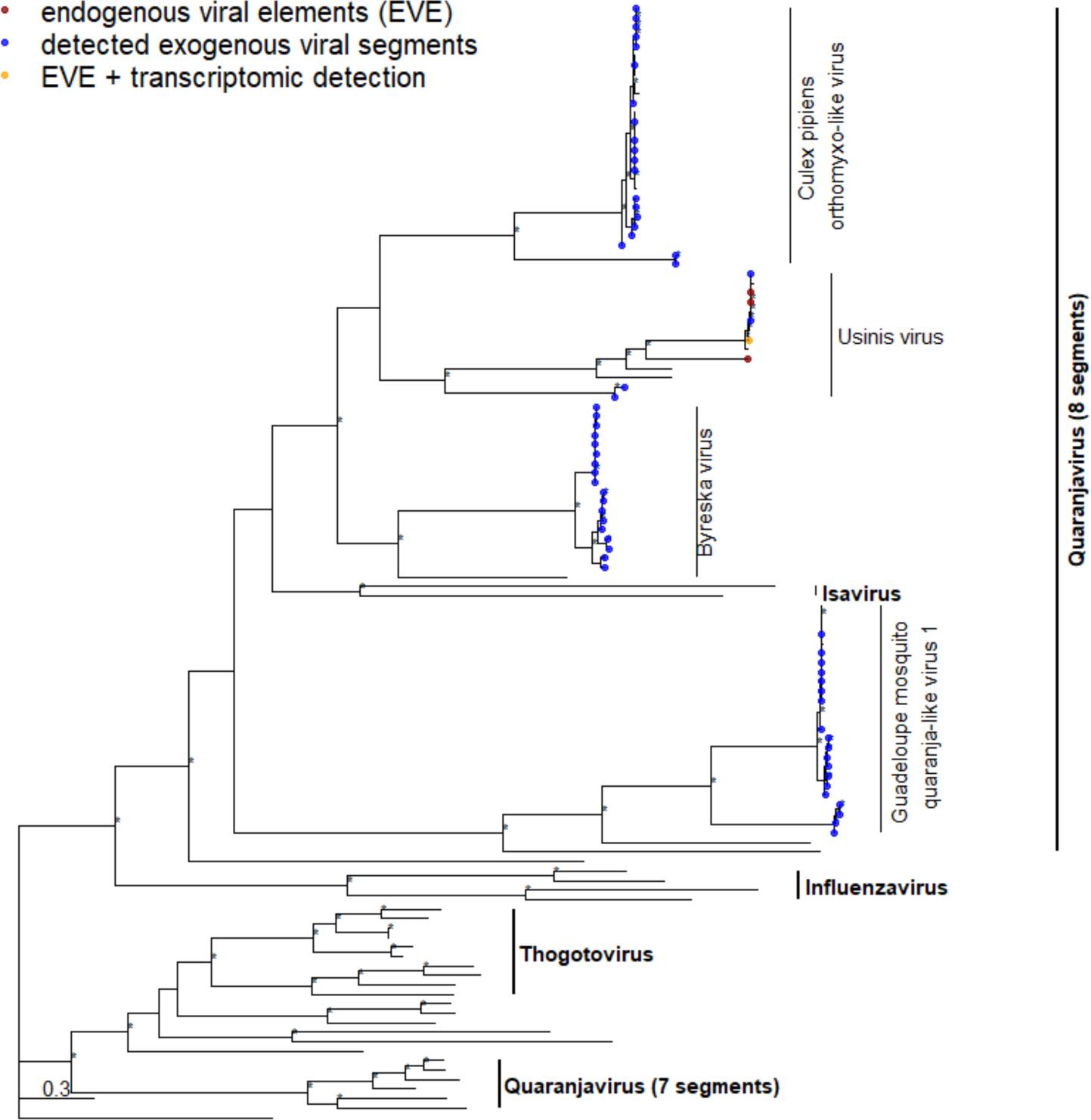
Phylogenetic tree depicting relationships between GP sequences of orthomyxoviruses. The maximum likelihood tree was constructed with model VT+I+G4 in IQ-TREE 1.6.12 (+ ultrafast bootstrap (1,000 replicates)). Significant ultrafast bootstrap values >95% are depicted with an asterisk. Detected protein sequences are marked in blue. Species and genus (bold) characteristics per clade are highlighted with black vertical lines to the right of the tree.

**Supplementary Figure 7:**
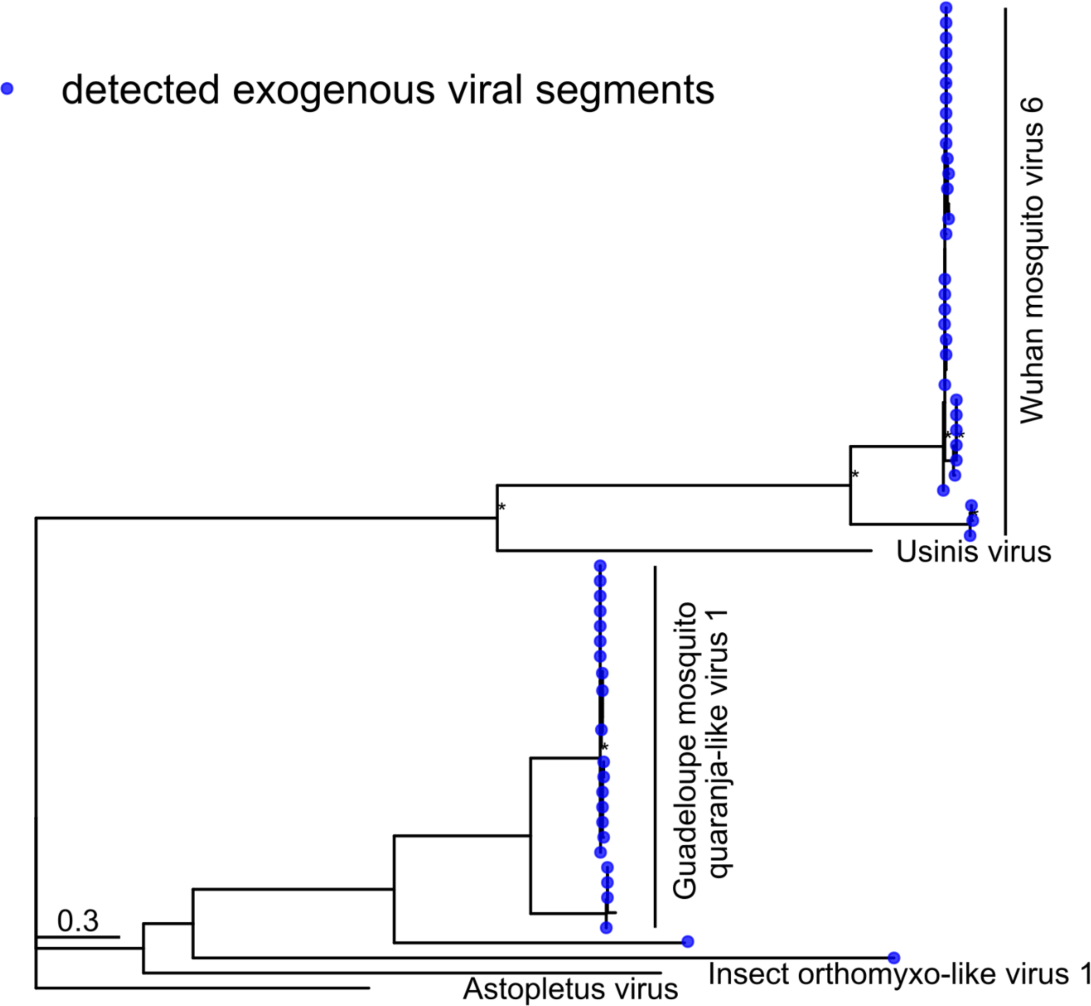
Phylogenetic tree depicting relationships between HP1 sequences of orthomyxoviruses. The maximum likelihood tree was constructed with model VT+G4 in IQ-TREE 1.6.12 (+ ultrafast bootstrap (1,000 replicates)). Significant ultrafast bootstrap values >95% are depicted with an asterisk. Detected protein sequences are marked in blue. Species and genus (bold) characteristics per clade are highlighted with black vertical lines to the right of the tree.

**Supplementary Figure 8:**
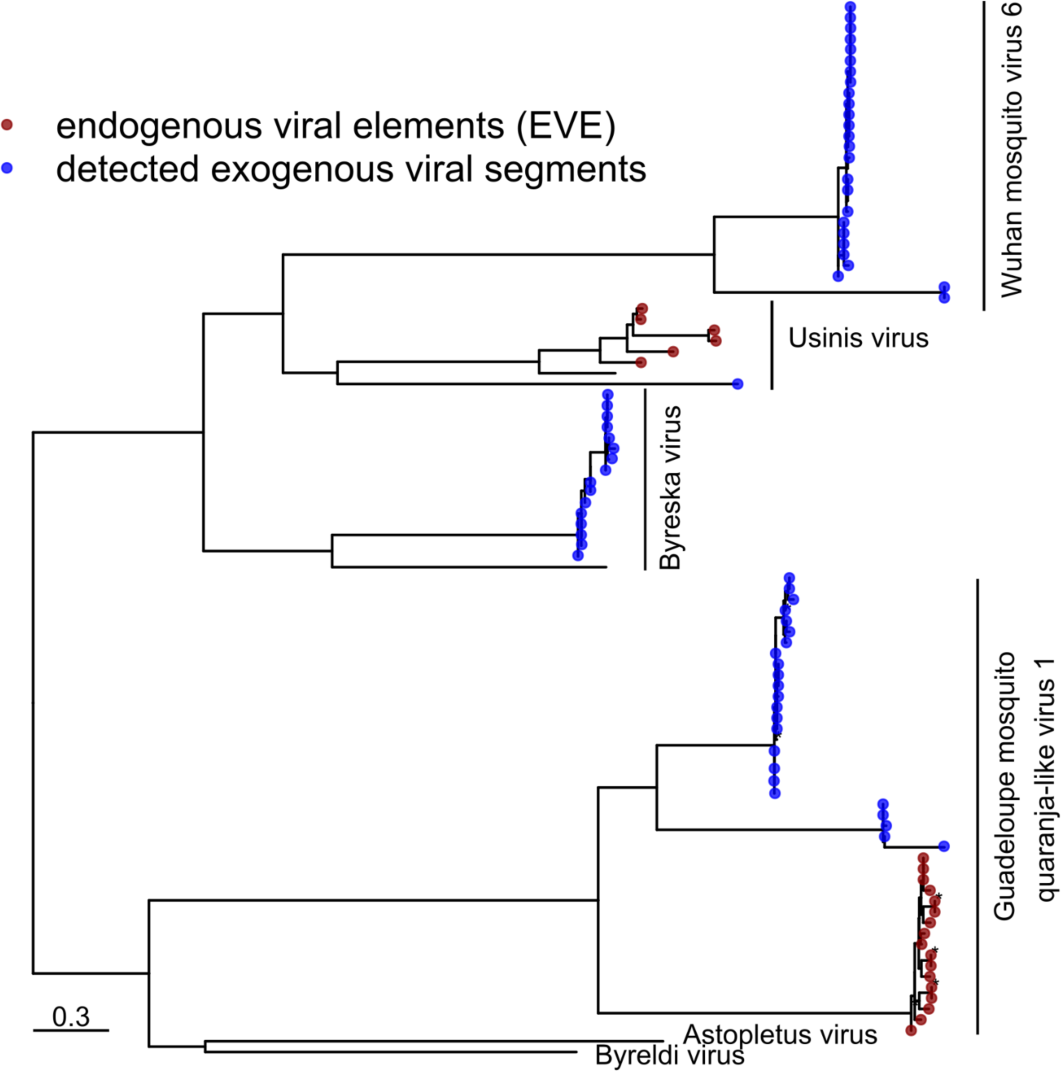
Phylogenetic tree depicting relationships between HP2 sequences of orthomyxoviruses. The maximum likelihood tree was constructed with model FLU+F+G4 in IQ-TREE 1.6.12 (+ ultrafast bootstrap (1,000 replicates)). Significant ultrafast bootstrap values >95% are depicted with an asterisk. Detected protein sequences are marked in blue. Species and genus (bold) characteristics per clade are highlighted with black vertical lines to the right of the tree.

**Supplementary Figure 9:**
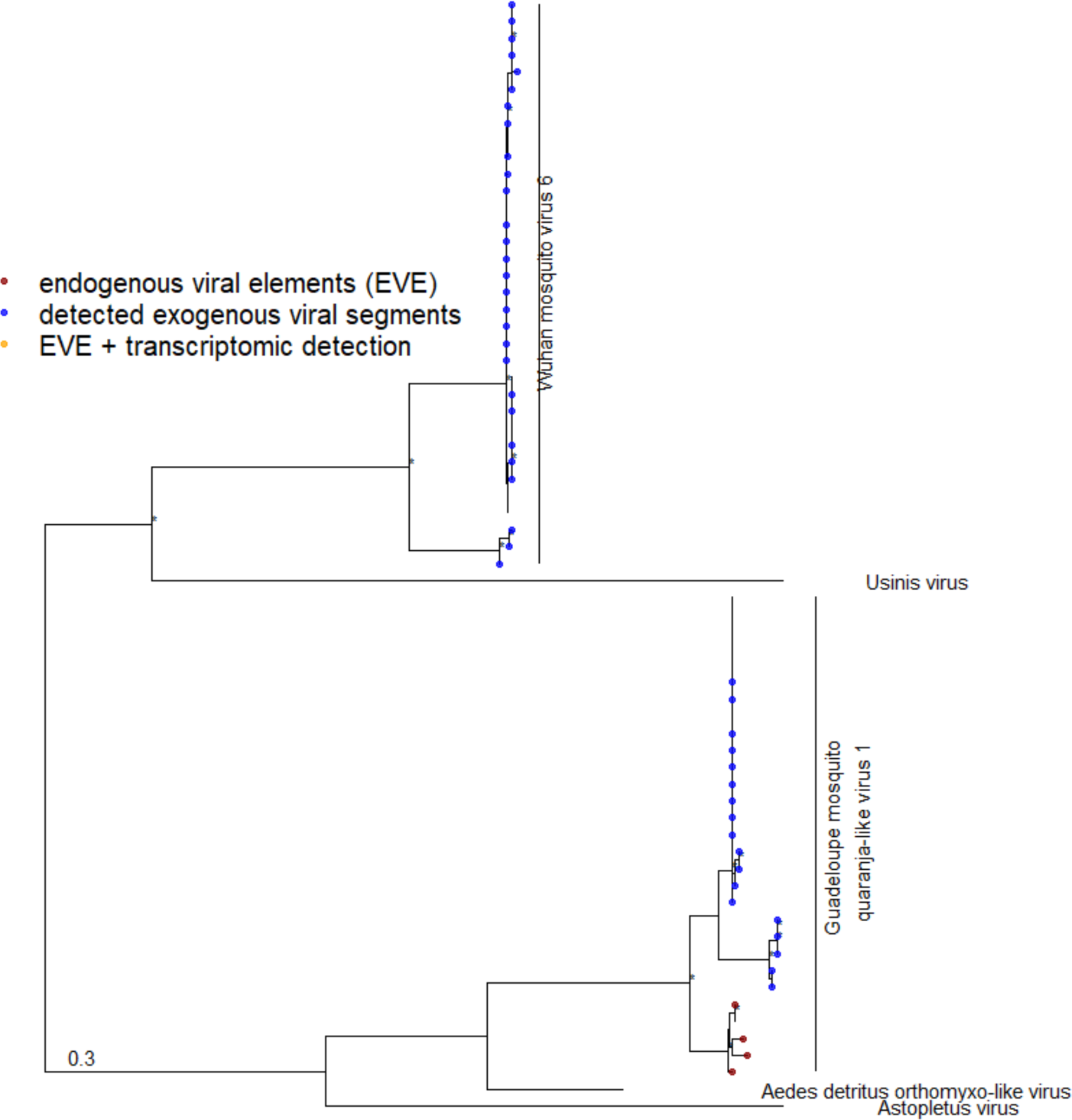
Phylogenetic tree depicting relationships between HP3 sequences of orthomyxoviruses. The maximum likelihood tree was constructed with model VT+I+G4 in IQ-TREE 1.6.12 (+ ultrafast bootstrap (1,000 replicates)). Detected protein sequences are marked in blue. Significant ultrafast bootstrap values >95% are depicted with an asterisk. Species and genus (bold) characteristics per clade are highlighted with black vertical lines to the right of the tree.

**Supplementary Figure 10:**
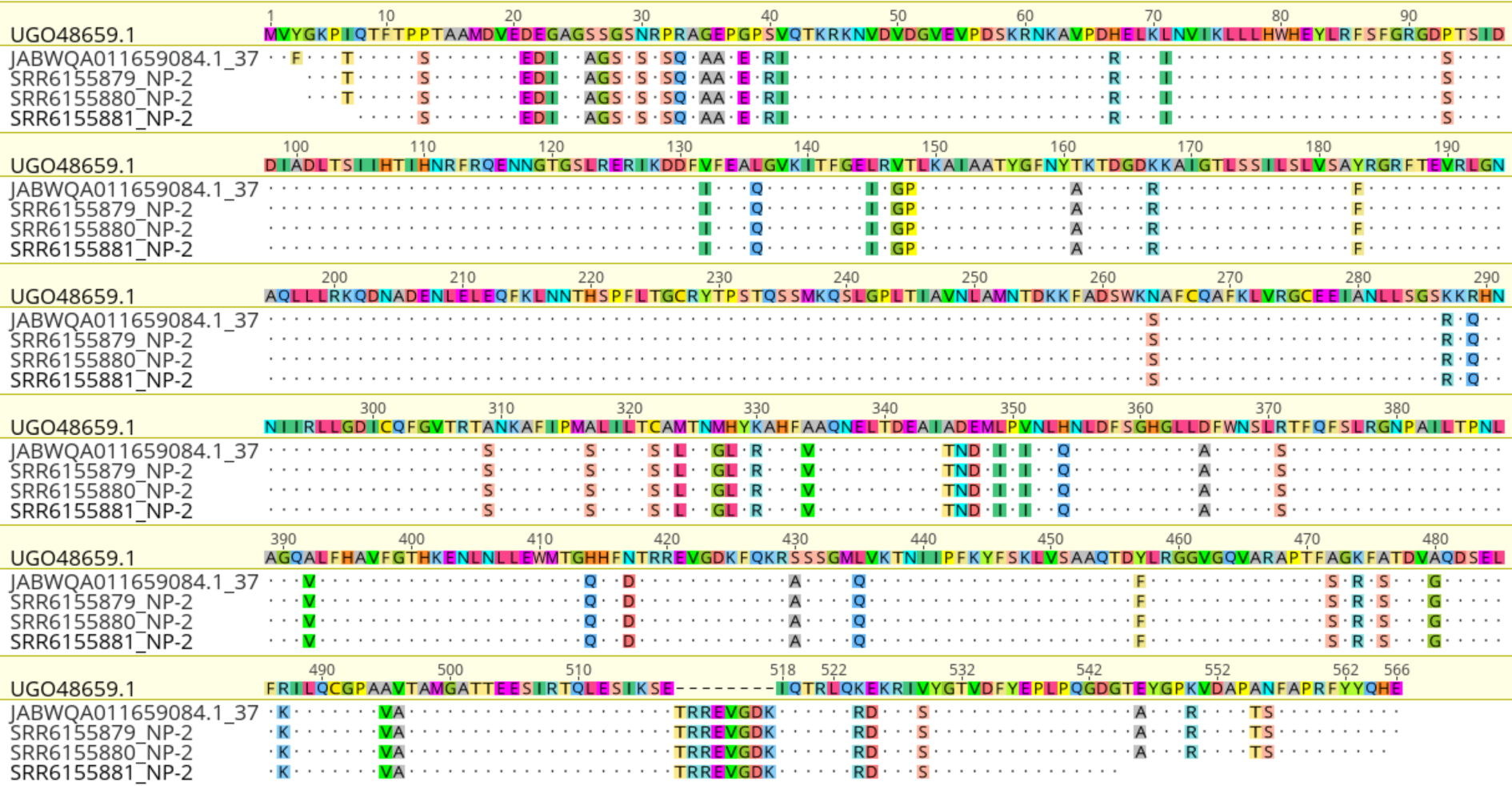
Sequence alignment of putative transcriptomic EVEs with full ORF.

**Supplementary Figure 11:**
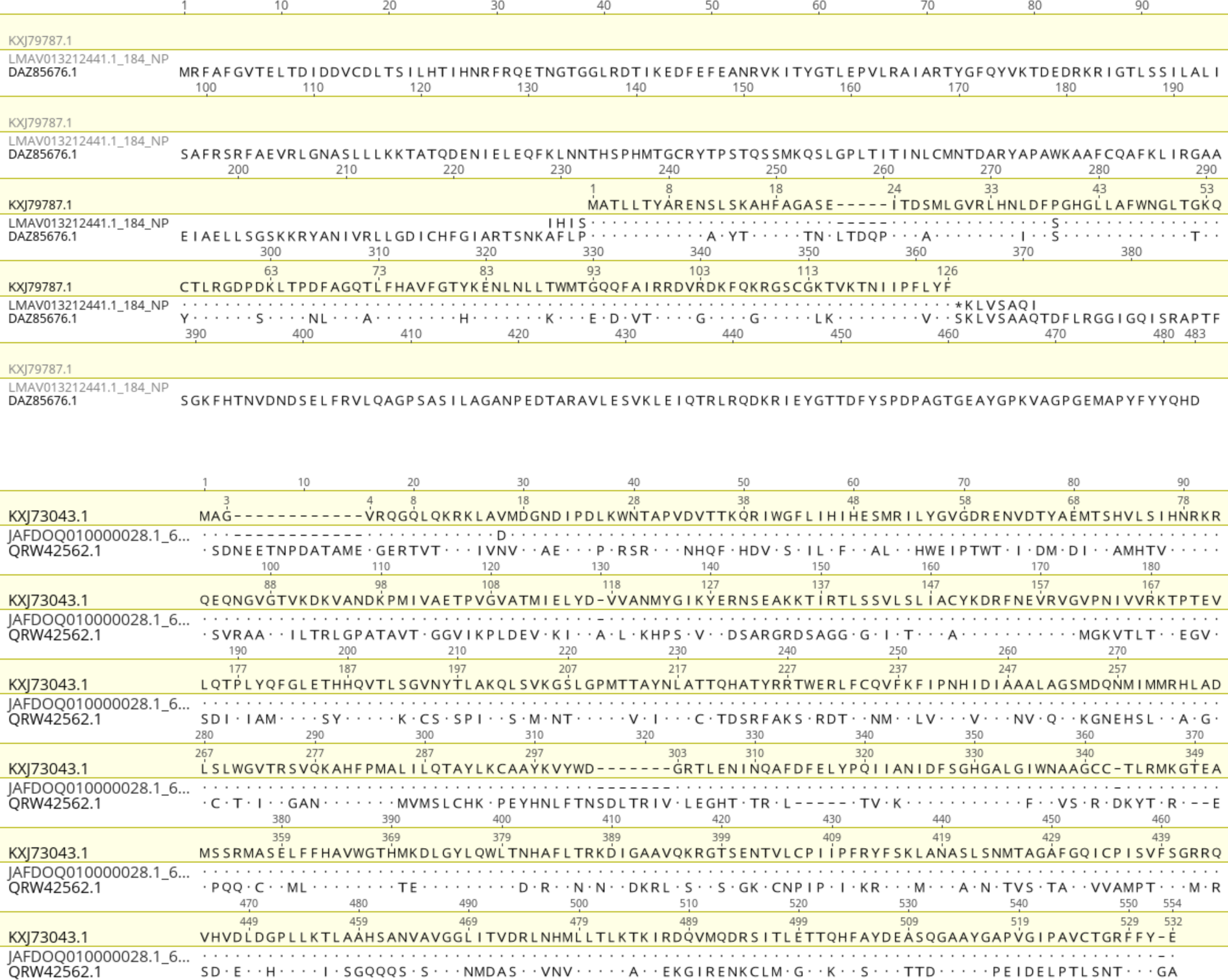
Sequence alignments between host hypothetical proteins and detected genomic EVE sequences.

## Notes

https://github.com/nbrait/EVEs-bias-virus-discovery

### Competing Interest Statement

The authors have declared no competing interest.

