## Supplementary Figure for "A tale of caution: How endogenous viral elements affect virus discovery in transcriptomic data"

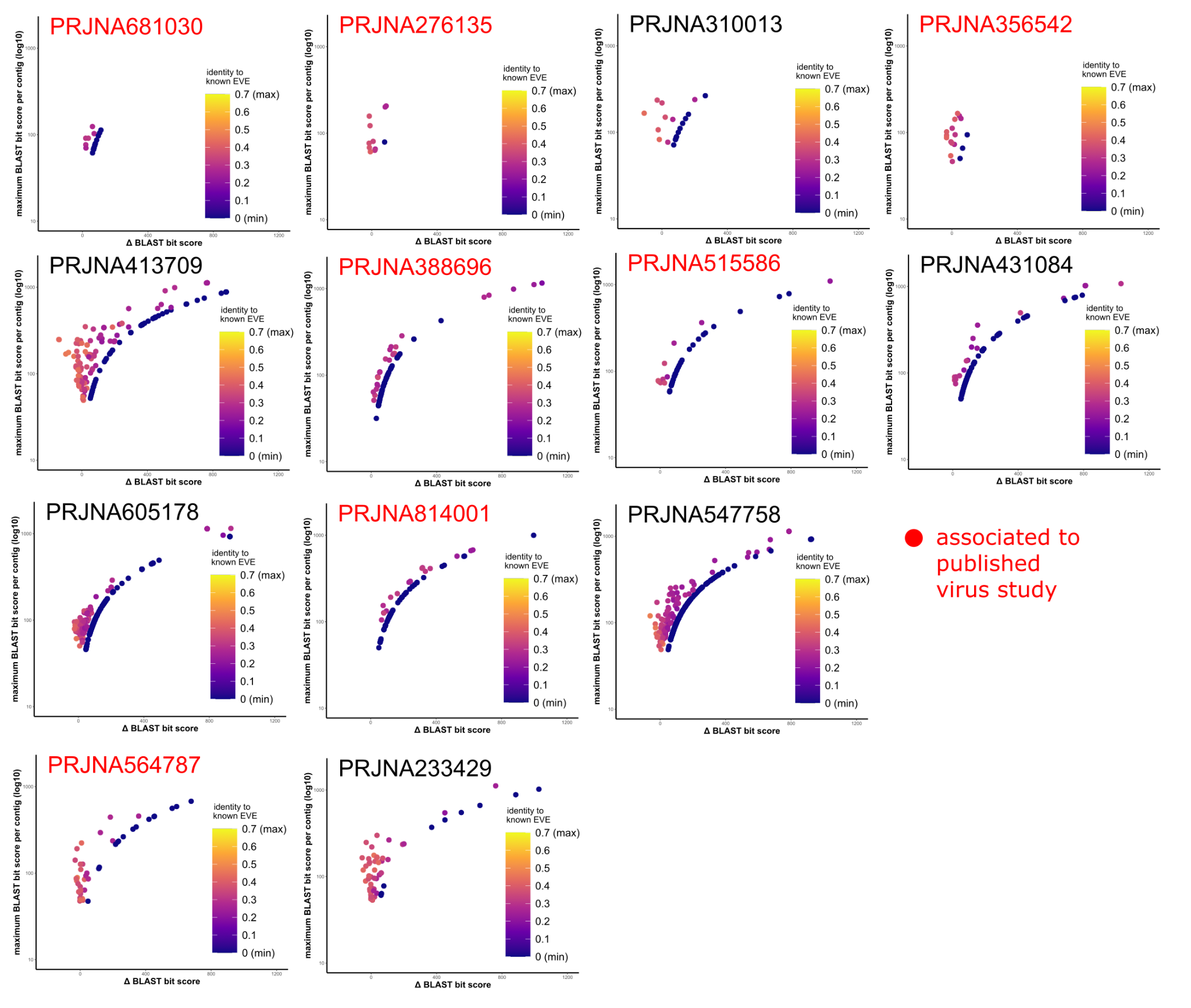

**Supplementary Figure 1: Plotted contig bit scores per bioproject**

Contigs were blasted against two custom databases: the existing *Orthomyxovirus* reference list or the translated amino-acid sequences of previously detected genomic EVEs. Data points are coloured according to aa sequence similarity to EVE references. Bioprojects associated with published virus studies are indicated in red. R script and details to bit score calculations and frameshift corrections can be found in Supplementary table 11.

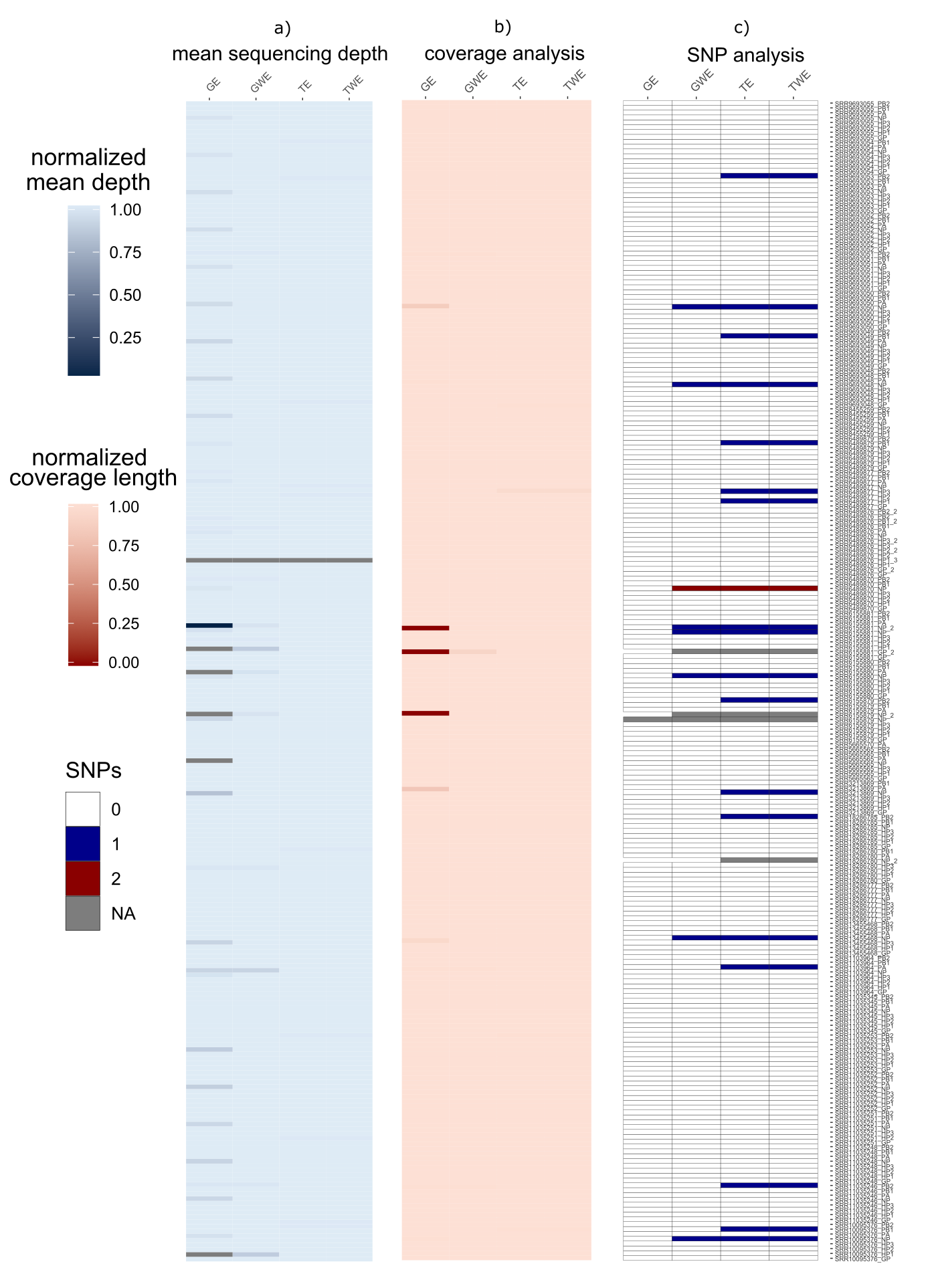

**Supplementary Figure 2 Comparison of mean sequence depths, coverage, and SNPs**

Sequence reads from four different pipelines were mapped back to the chimeric reference sequences generated in Geneious Prime. Sequencing depths were generated with samtools. The percentages of length coverage were calculated by dividing the corrected coverage lengths minus the count of Ns by full coverage length. For SNP calculations, GE (genomic host read removal with EVE presence) was used as a reference sequence to observe differences in the other three workflows. a) Mean sequencing depth analysis, b) coverage analysis, c) SNP analysis. Abbreviations: GE (genomic host read removal with EVE presence), GWE (genomic host read removal without EVE presence), TE (transcriptomic host read removal with EVE presence), TWE (transcriptomic host read removal without EVE presence). Segments that did not generate sam output files after bowtie2 mapping are depicted in gray.

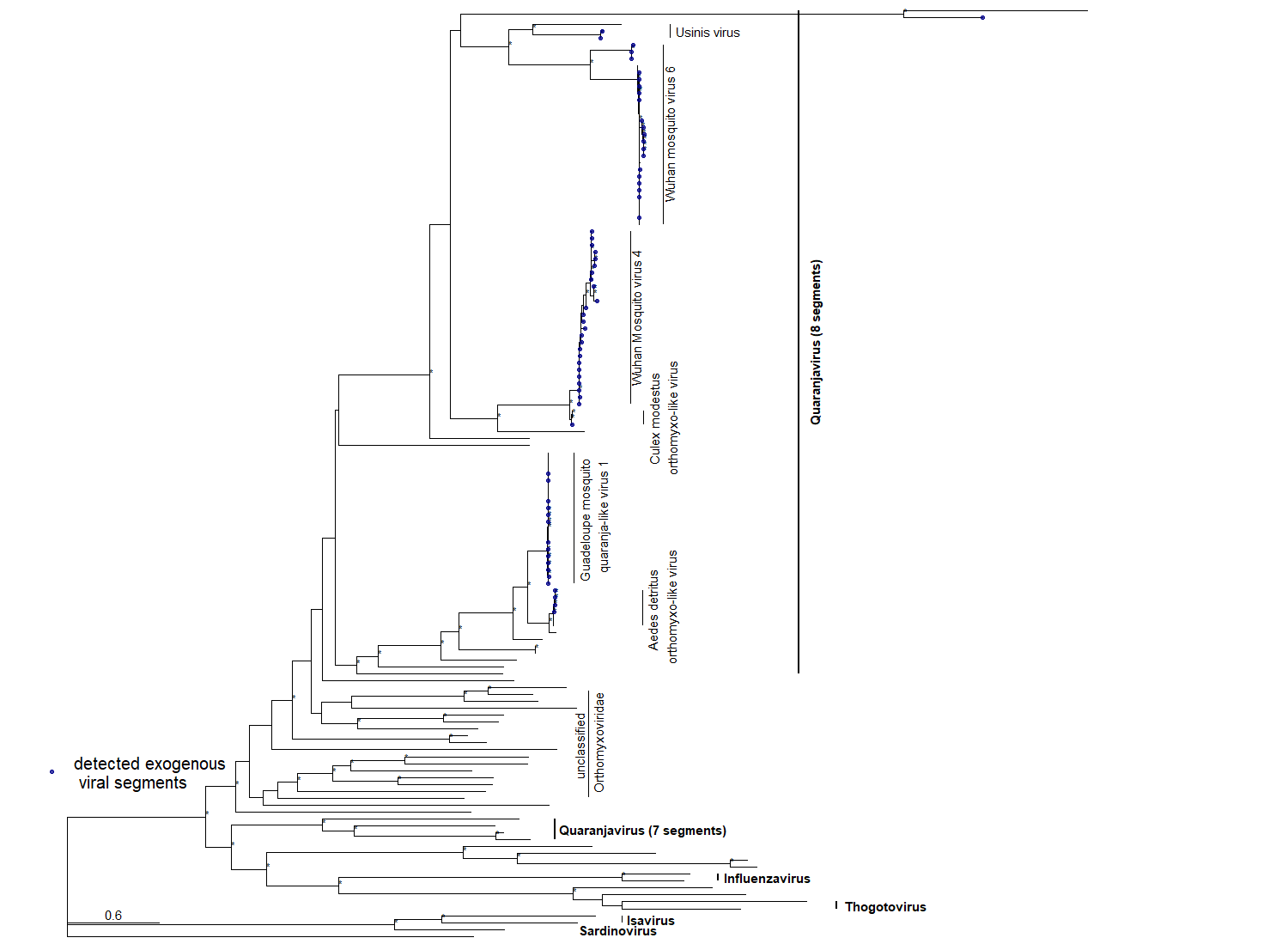

**Supplementary Figure 3: Phylogenetic tree depicting relationships between PB2 sequences of orthomyxoviruses**. The maximum likelihood tree was constructed with model VT+I+G4 in IQ-TREE 1.6.12 (+ ultrafast bootstrap (1,000 replicates)). Significant ultrafast bootstrap values >95% are depicted with an asterisk. Detected protein sequences are marked in blue. Species and genus (bold) characteristics per clade are highlighted with black vertical lines to the right of the tree.

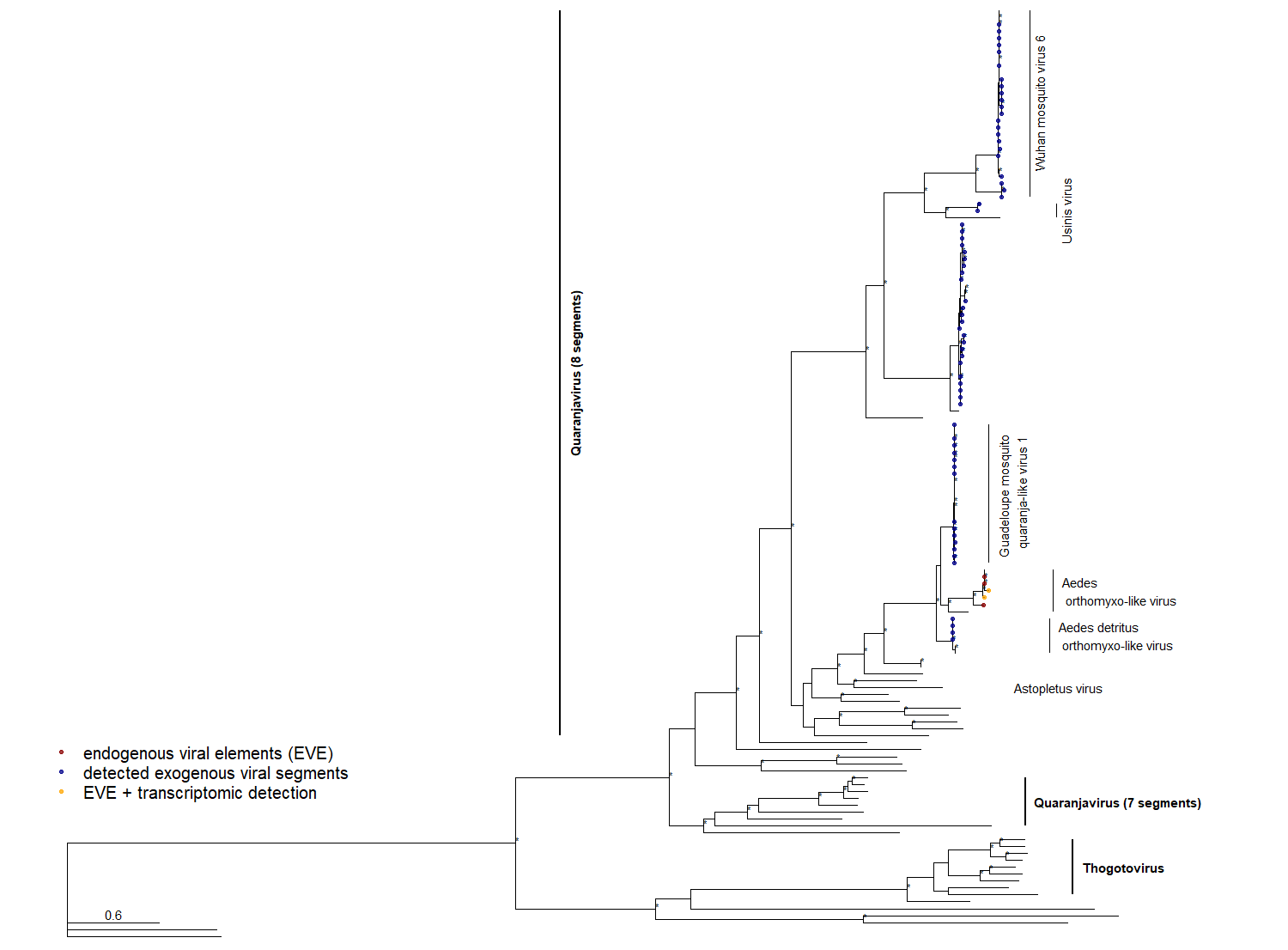

**Supplementary Figure 4: Phylogenetic tree depicting relationships between PB1 sequences of orthomyxoviruses**. The maximum likelihood tree was constructed with model VT+I+G4 in IQ-TREE 1.6.12 (+ ultrafast bootstrap (1,000 replicates)). Significant ultrafast bootstrap values >95% are depicted with an asterisk. Detected protein sequences are marked in blue. Species and genus (bold) characteristics per clade are highlighted with black vertical lines to the right of the tree.

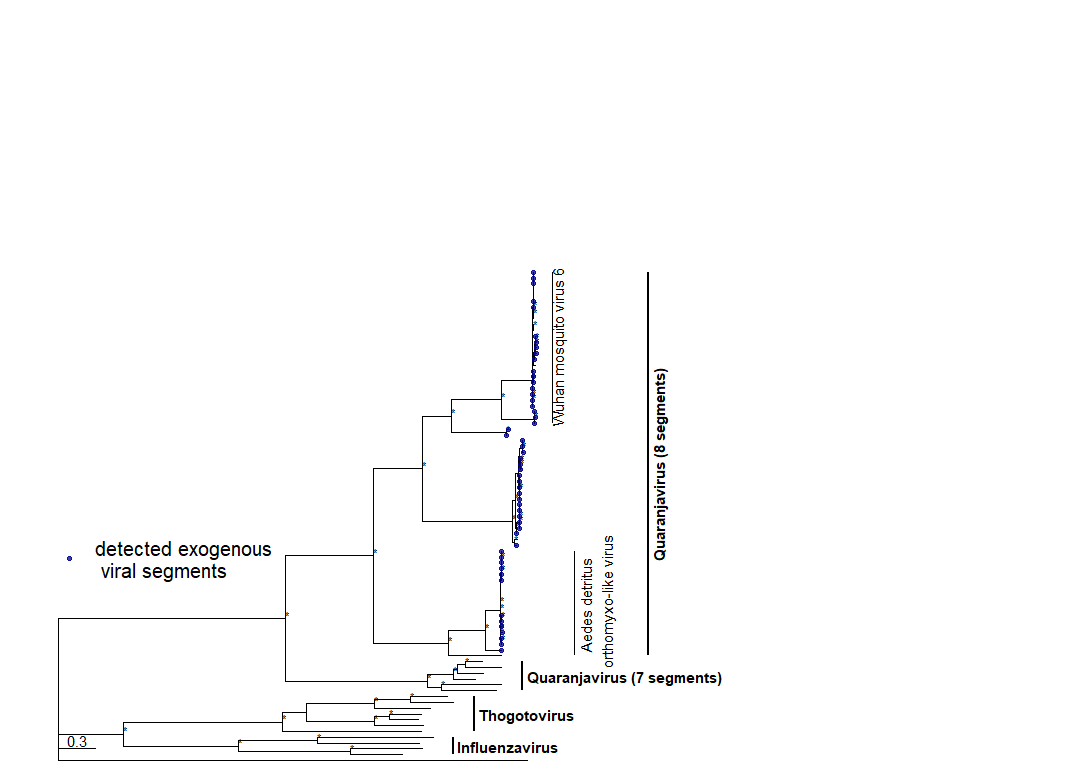

**Supplementary Figure 5: Phylogenetic tree depicting relationships between PA sequences of orthomyxoviruses.** The maximum likelihood tree was constructed with model VT+I+G4 in IQ-TREE 1.6.12 (+ ultrafast bootstrap (1,000 replicates)). Significant ultrafast bootstrap values >95% are depicted with an asterisk. Detected protein sequences are marked in blue. Species and genus (bold) characteristics per clade are highlighted with black vertical lines to the right of the tree.

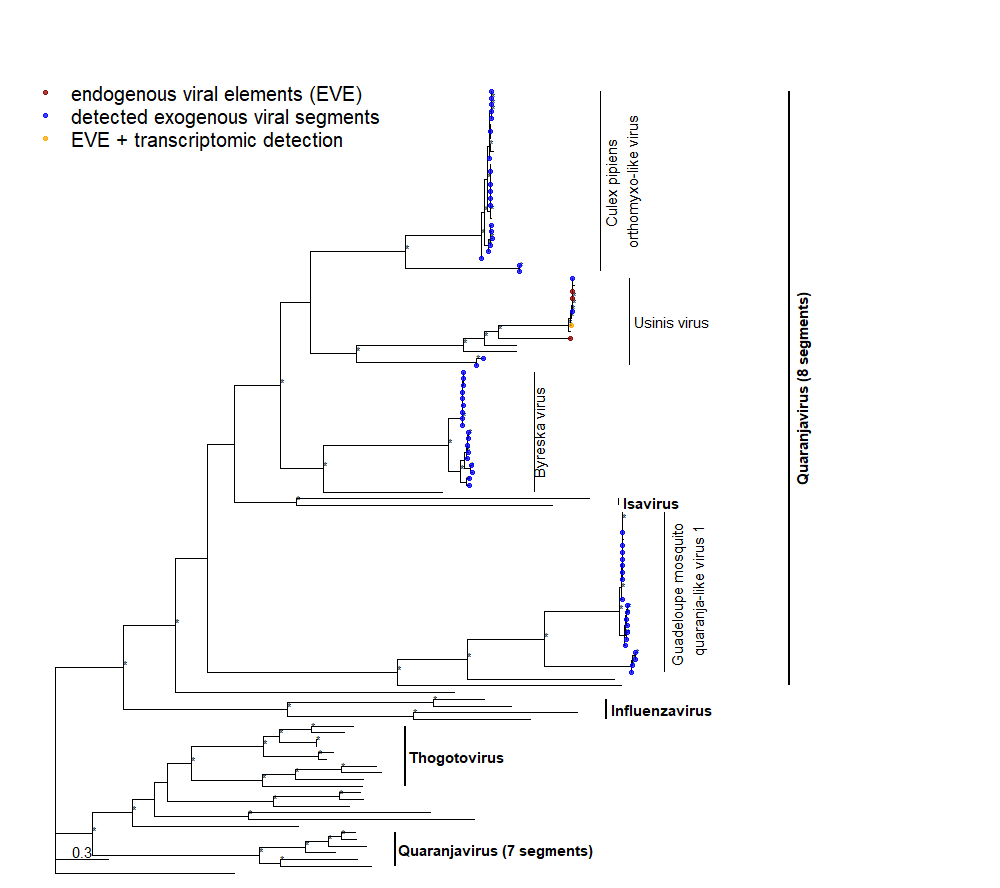

**Supplementary Figure 6:** **Phylogenetic tree depicting relationships between GP sequences of orthomyxoviruses**. The maximum likelihood tree was constructed with model VT+I+G4 in IQ-TREE 1.6.12 (+ ultrafast bootstrap (1,000 replicates)). Significant ultrafast bootstrap values >95% are depicted with an asterisk. Detected protein sequences are marked in blue. Species and genus (bold) characteristics per clade are highlighted with black vertical lines to the right of the tree.

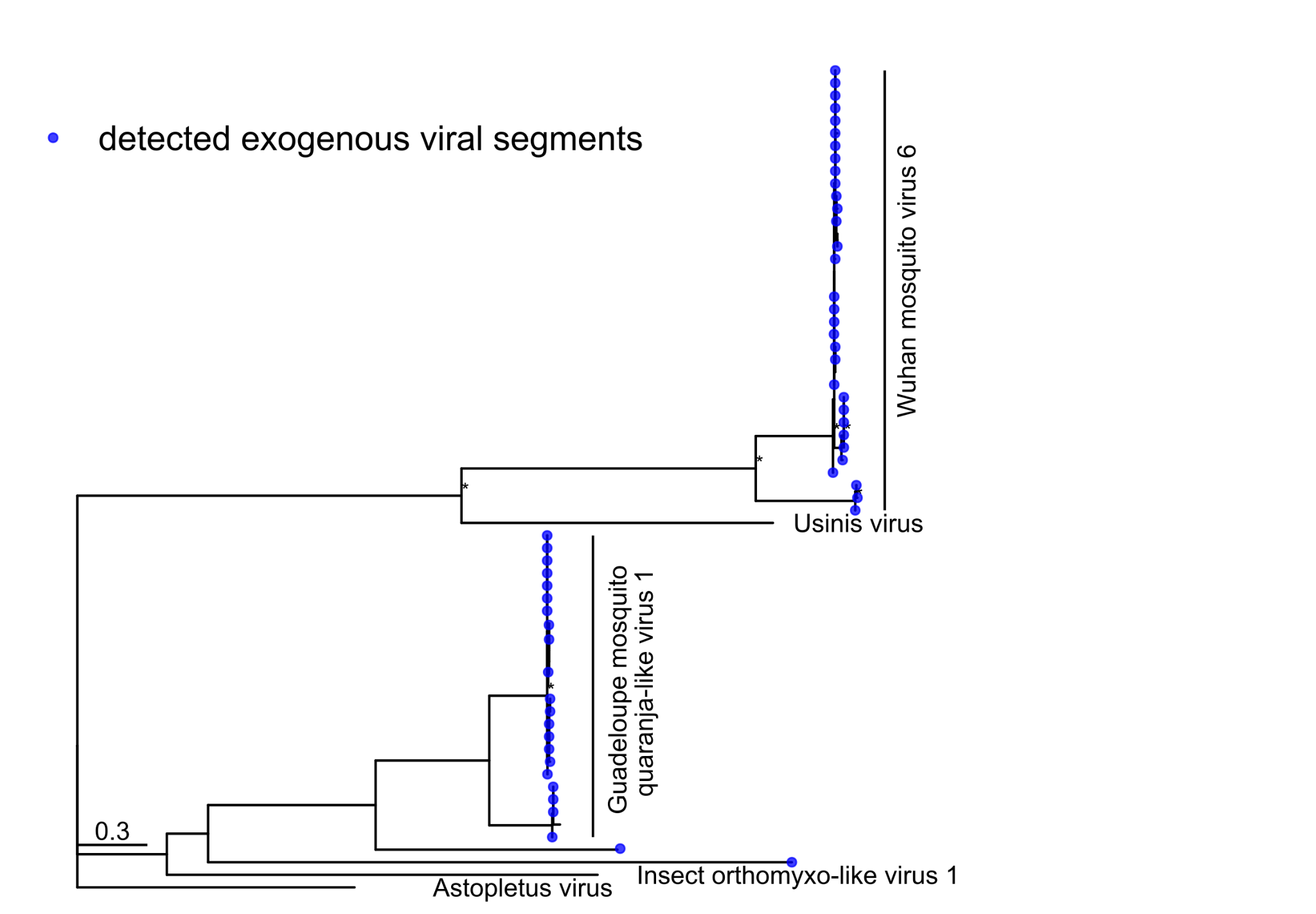

**Supplementary Figure 7:** **Phylogenetic tree depicting relationships between HP1 sequences of orthomyxoviruses**. The maximum likelihood tree was constructed with model VT+G4 in IQ-TREE 1.6.12 (+ ultrafast bootstrap (1,000 replicates)). Significant ultrafast bootstrap values >95% are depicted with an asterisk. Detected protein sequences are marked in blue. Species and genus (bold) characteristics per clade are highlighted with black vertical lines to the right of the tree.

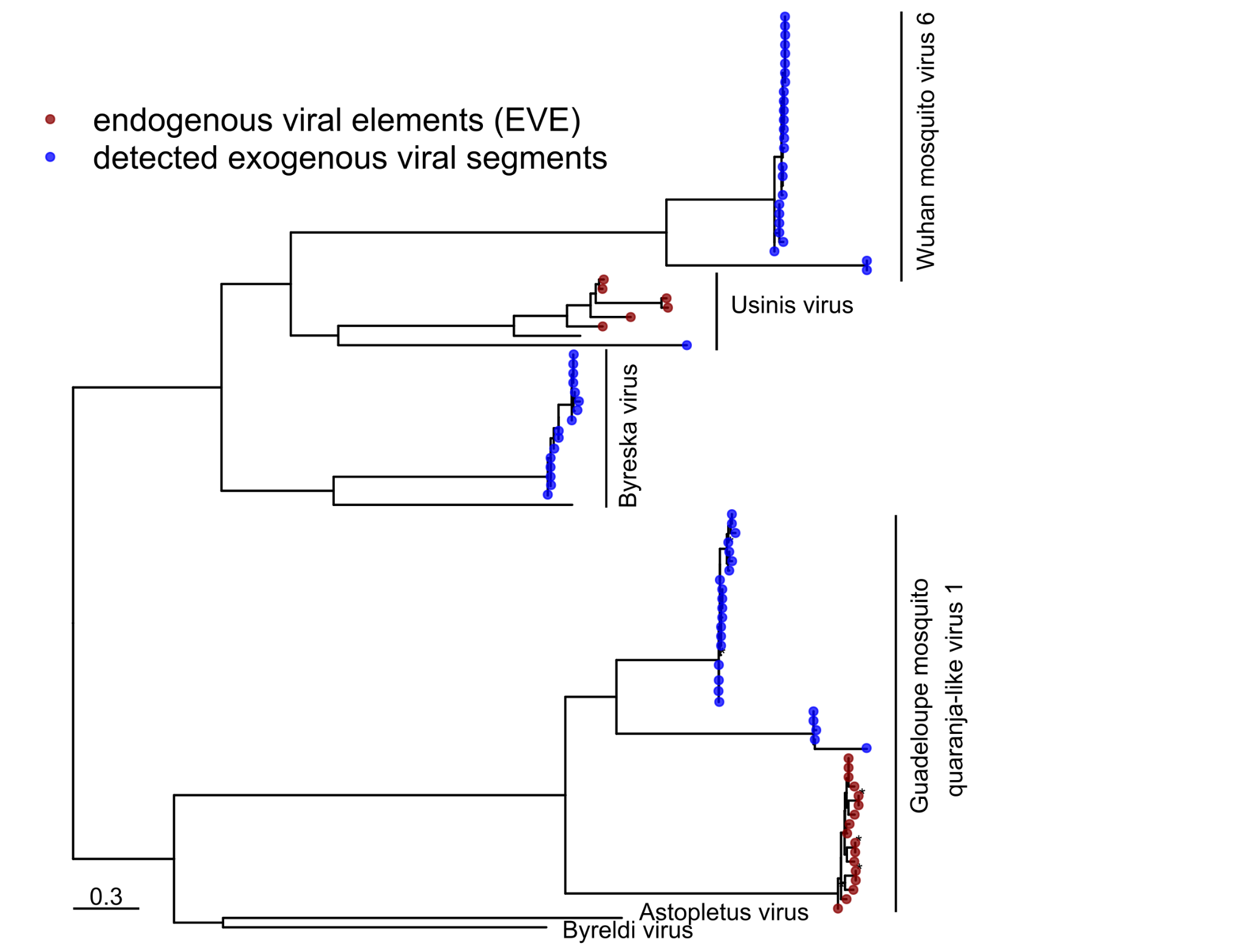

**Supplementary Figure 8:** **Phylogenetic tree depicting relationships between HP2 sequences of orthomyxoviruses**. The maximum likelihood tree was constructed with model FLU+F+G4 in IQ-TREE 1.6.12 (+ ultrafast bootstrap (1,000 replicates)). Significant ultrafast bootstrap values >95% are depicted with an asterisk. Detected protein sequences are marked in blue. Species and genus (bold) characteristics per clade are highlighted with black vertical lines to the right of the tree.

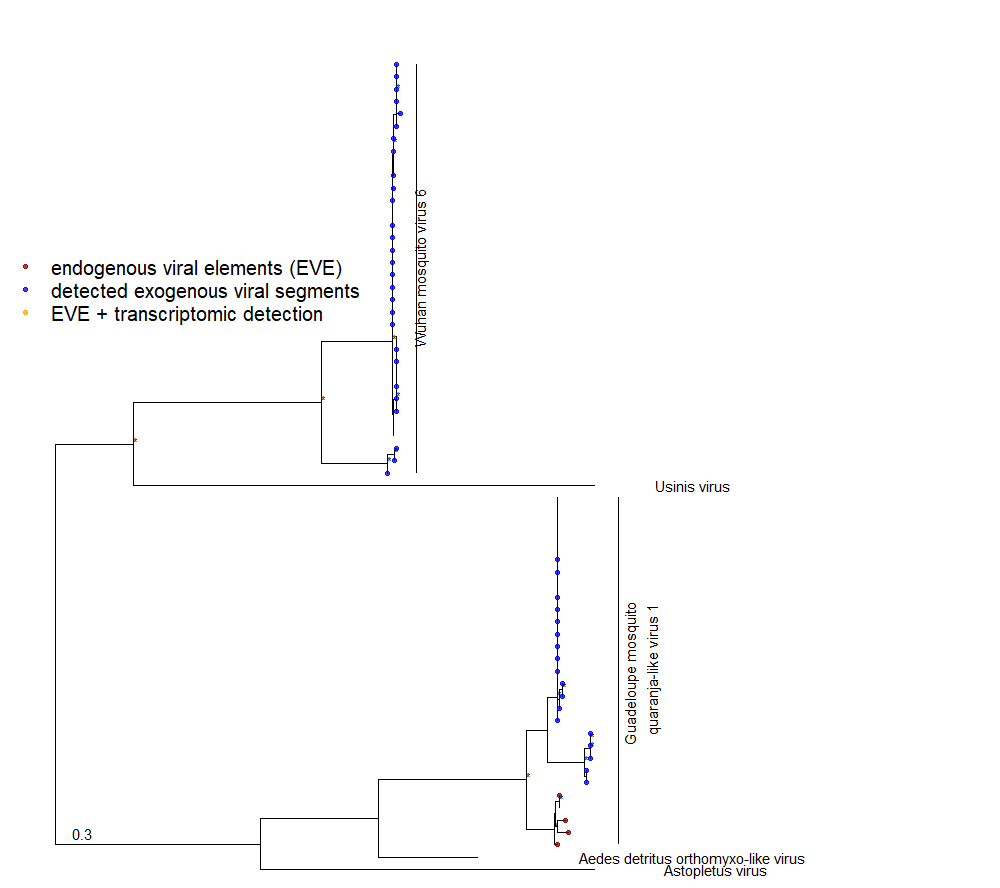

**Supplementary Figure 9:** **Phylogenetic tree depicting relationships between HP3 sequences of orthomyxoviruses**. The maximum likelihood tree was constructed with model VT+I+G4 in IQ-TREE 1.6.12 (+ ultrafast bootstrap (1,000 replicates)). Detected protein sequences are marked in blue. Significant ultrafast bootstrap values >95% are depicted with an asterisk. Species and genus (bold) characteristics per clade are highlighted with black vertical lines to the right of the tree.

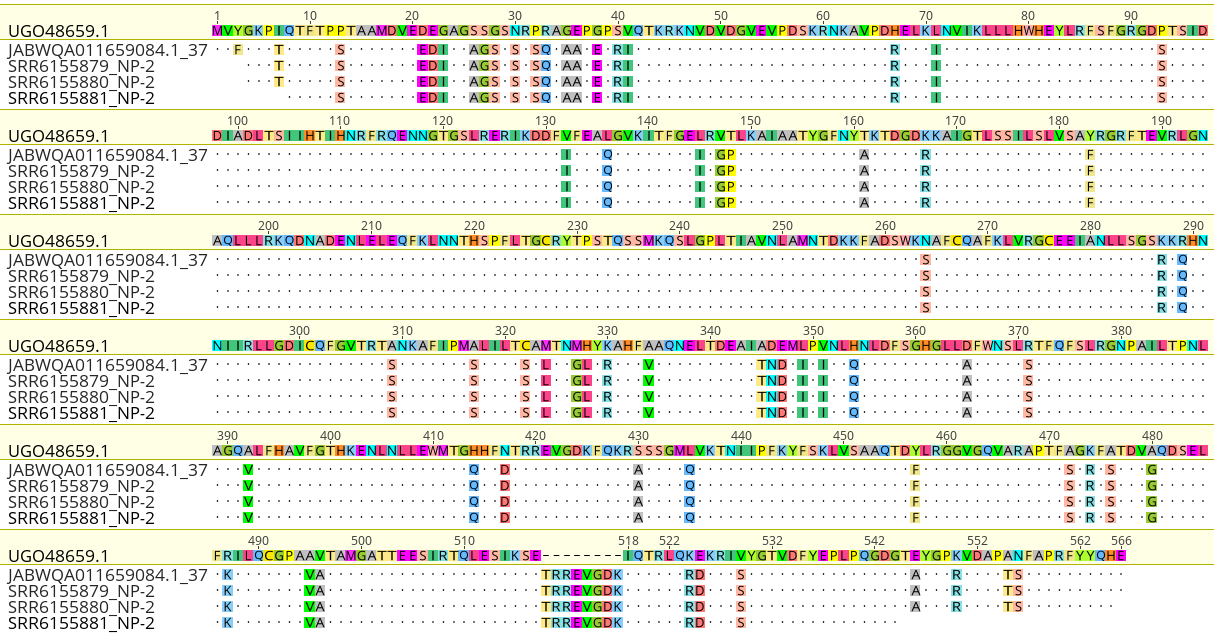

**Supplementary Figure 10: Sequence alignment of putative transcriptomic EVEs with full ORF**

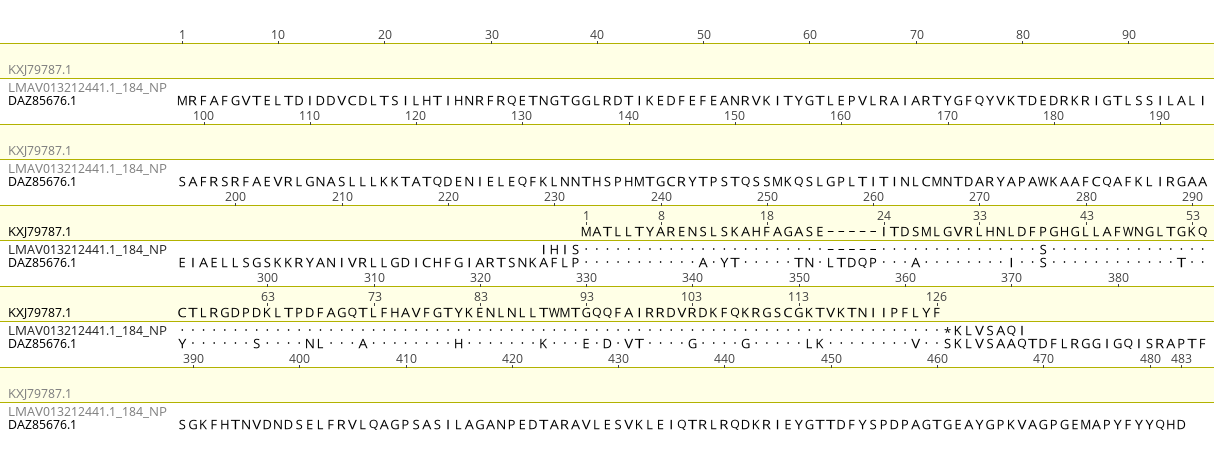

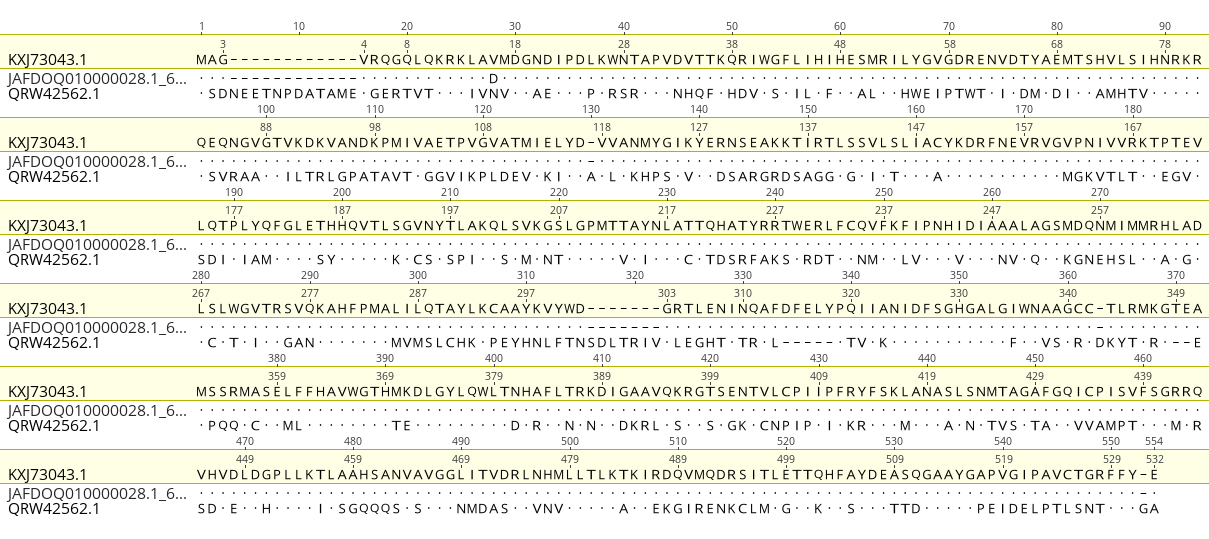

**Supplementary Figure 11: Sequence alignments between host hypothetical proteins and detected genomic EVE sequences**
